# Wetness modulates the effects of grazing on net ecosystem productivity in global grasslands

**DOI:** 10.64898/2026.01.27.701613

**Authors:** Yueqiang Wu, Le Qi, Hao Li, Jiguang Feng, Peng Zhou, Hangyu Li, Xiaoyang Gao, Zhijie Wang, Shilin Cui, Ping Yin, Wenhong Ma, Cunzhu Liang, Zhiyong Li, Biao Zhu

## Abstract

Overgrazing has induced widespread grassland degradation globally. However, how grazing intensity affects ecosystem carbon dioxide (CO_2_) fluxes with wetness fluctuations in grasslands worldwide remains poorly understood. Here we measured ecosystem CO_2_ fluxes covering the 7th to 11th years of a continuous grazing experiment in a typical steppe, and conducted a meta-analysis of 585 observations by collecting relevant data in global grasslands. Our experimental results showed that mean gross primary productivity (GPP) and ecosystem respiration (ER) were decreased under heavy grazing (HG) compared with control without grazing (CK), whereas net ecosystem productivity (NEP) was not different between HG and CK in the typical steppe. The NEP was higher under light grazing in the wetter years, but it was not different between light grazing and CK under drier years in the typical steppe. The structural equation model showed that grazing affected NEP by increasing plant relative growth rate, while decreasing plant aboveground biomass (AGB). In contrast, grazing decreased GPP, ER and NEP, respectively, in global grasslands. Both our field experiment and the meta-analysis revealed that the response of NEP to light grazing, rather than heavy grazing, was linearly correlated with the wetness index. Higher wetness and AGB increased the response of NEP to grazing in global grasslands. However, heavy grazing reduced NEP and AGB even under higher wetness indices in global grasslands, resulting from a loss of their resilience in long-term heavy grazing. These findings indicate that light grazing appears to be a promising management to promote plant relative growth rate and carbon sequestration. Overall, the meta-analysis and field experiment jointly provide global perspectives on the response of ecosystem CO_2_ fluxes to grazing intensity and improve our understanding of the factors influencing the response of ecosystem CO_2_ fluxes to grazing intensity.

## 1. Introduction

Globally, grassland ecosystems cover approximately one fifth of the world land surface and contain about 20-30% of the global pool of soil organic carbon (SOC) (Du et al., 2022). Grasslands also play a vital role in livestock production, promoting carbon (C) sequestration and regulating climate change (Heimann & Reichstein, 2008; Zhou et al., 2017). Grazing is a widespread land-use activity that alters terrestrial C dynamics (He et al., 2020; Zhou et al., 2025). However, grassland ecosystems have seriously deteriorated globally (Zhang et al., 2025). Grassland degradation is influenced by a multitude of driving factors, such as overgrazing and precipitation limitation (Sloat et al., 2018). Furthermore, the C balance of grasslands is also increasingly influenced by changes in grazing management practices and water availability (Hilker et al., 2014; Lemoine et al., 2016; Ren et al., 2025). Overgrazing and drier conditions could potentially drive grasslands from acting as C sinks to C sources (Hao et al., 2018; Morgan et al., 2016; Shao et al., 2013).

Gross primary productivity (GPP) and ecosystem respiration (ER) are critical processes governing C sequestration and emission in terrestrial ecosystems (Chen et al., 2023; Knauer et al., 2023). Net ecosystem productivity (NEP) is the balance between ecosystem C uptake and release, and serves as a key indicator of the net C exchange (Niu et al., 2010; Zhang et al., 2024). Grazing influences NEP by altering GPP and ER (Fig. 1). Grazing directly removes aboveground plant tissues through herbivory, altering vegetation cover, soil temperature, and aboveground biomass (AGB) and thereby influencing plant photosynthesis and GPP (Owensby et al., 2006; Zhou et al., 2007). Plants with high relative growth capacity may rapidly restore leaf area and photosynthesis tissues after grazing, potentially maintaining or even enhancing GPP (Oesterheld & McNaughton, 1991; Penner & Frank, 2021). In contrast, heavy grazing or water limitation may constrain plant regrowth, limit soil moisture, leaf area recovery, and reduce GPP. Grazing may also affect ER by reducing litter and root carbon inputs, altering belowground carbon allocation, and modifying soil bulk density, porosity, water infiltration, and aeration through livestock tamping, with consequences for root respiration and microbial decomposition (Bardgett & Wardle, 2003; Wu et al., 2023; Xun et al., 2018). Variation in NEP responses to grazing may depend on whether post-grazing recovery of GPP can compensate for carbon losses through ER. Thus, plant relative growth rate and compensatory growth capacity may represent important mechanistic traits linking grazing disturbance, photosynthesis recovery, GPP dynamics, and NEP response.

**Fig. 1.**
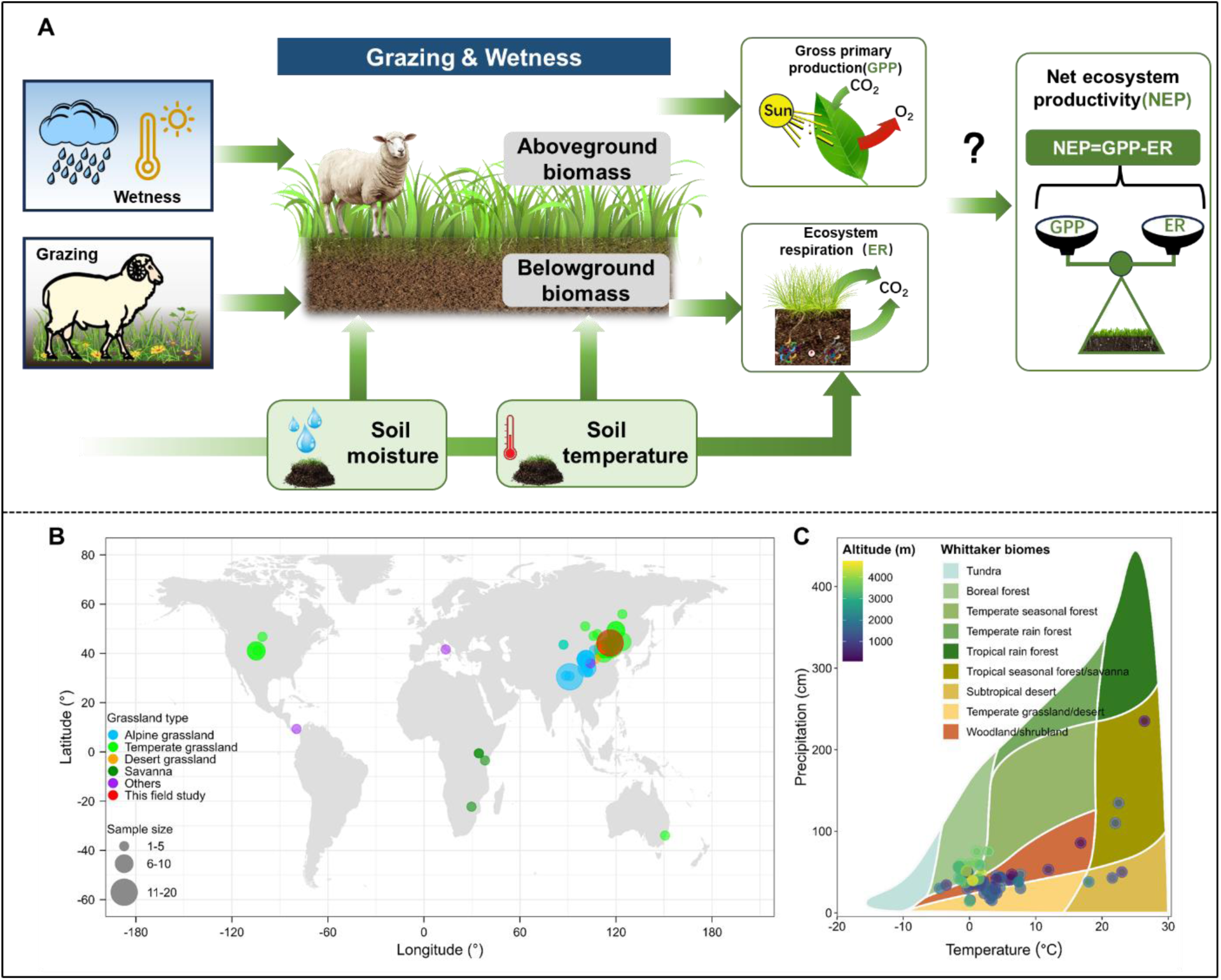
Conceptual illustration of how grazing with wetness fluctuation would affect net ecosystem productivity (A), and geographic (B) and climatic distributions of data (C) included in this meta-analysis.

Assessing ecosystem carbon dioxide (CO_2_) fluxes (GPP, ER and NEP) in grasslands is essential to the terrestrial C cycle at local, regional and global scales. However, current evidence on the effect of grazing on ecosystem CO_2_ fluxes in grasslands remains inconsistent, with studies reporting both positive (Chang et al., 2020; Du et al., 2022) and negative (Liu et al., 2020; Wang et al., 2023; Zhu et al., 2015) effects. For instance, some studies demonstrated that grazing decreased GPP and ER (Nakano et al., 2020; Qi et al., 2024; Wang et al., 2025), while others reported an increase or no difference in GPP or ER (Chang et al., 2020; Luo et al., 2015; Shi et al., 2022). Similarly, some studies investigated that grazing significantly reduced NEP (Wang et al., 2023; Zhu et al., 2015), while others reported an increase or no difference in NEP under grazing (Du et al., 2022; Gomez-Casanovas et al., 2018; Yu et al., 2025). The variations in the responses of ecosystem CO_2_ fluxes are likely driven by differences in grazing intensities, climatic variations, livestock and grassland types (Hou et al., 2016; Wang et al., 2023; Xiao et al., 2011; Zhu et al., 2015).

The effects of grazing intensities on ecosystem CO_2_ fluxes in global grasslands were also varied in different studies (Table S1; Jiang et al., 2020; Rong et al., 2017; Zhang et al., 2022). Specifically, heavy grazing usually reduced the GPP, ER and NEP (Ju et al., 2024; Rong et al., 2017; Zhang et al., 2022), but a meta-analysis also reported the response of NEP was increased in Chinese grasslands (Jiang et al., 2020). Generally, light grazing can enhance NEP by promoting plant regeneration and improving photosynthetic efficiency, whereas moderate to heavy grazing tends to reduce NEP, largely as a result of decreased vegetation coverage and plant biomass (Yu et al., 2025). Conversely, a meta-analysis showed that the response of NEP was decreased under light and moderate grazing (Zhang et al., 2022). However, it was reported that moderate grazing led to the highest NEP (Zhu et al., 2015). These conflicting results underscore the complex responses of ecosystem CO_2_ fluxes in grasslands to grazing and highlight the necessity to identify key factors controlling ecosystem CO_2_ flux dynamics under grazing management. This understanding is essential for accurately modelling and predicting grassland C dynamics under global change.

Many studies have demonstrated that interannual variation in grassland NEP can be substantial, primarily driven by fluctuating climatic conditions, as they can directly regulate vegetation growth via altering soil moisture and temperature (Polley et al., 2008; Rong et al., 2017; Shen et al., 2016). Annual precipitation is one of the climatic parameters, while the wetness index (WI) serves as a more integrative climatic indicator that incorporates both precipitation and temperature, thereby reflecting the overall water surplus or deficit (Song et al., 2019). A higher wetness index (WI > 30) indicates sufficient water availability for plant growth, whereas a lower wetness index (WI ≤ 30) suggests the water availability may be limited (De Martonne, 1926). Considerable uncertainty remains regarding how NEP responds to grazing under different water conditions. Although evidence suggested that, under wet year, grazing significantly increased NEP; under dry year, grazing suppressed NEP and can even turn NEP negative (Shao et al., 2013). In general, grazing tends to have a less suppressive effect on NEP under wet than dry conditions (Okach et al., 2019; Zhou et al., 2019). For example, heavy grazing consistently reduced NEP, with a more pronounced decline occurring during dry years in a desert steppe (Jin et al., 2023). In addition, overgrazing led to decoupling of precipitation patterns and ecosystem carbon exchange in the desert steppe, primarily by altering plant community composition (Wang et al., 2023). However, it remains unclear how NEP responds to grazing intensities under water availability fluctuations across global grasslands.

Previous studies on the effects of grazing intensities on ecosystem CO_2_ fluxes have been constrained by limited spatial and temporal scales, which has led to an incomplete understanding of how different grazing intensities influence ecosystem CO_2_ fluxes. Furthermore, does the wetness modulate the effect of grazing on ecosystem CO_2_ fluxes in the typical steppe? Are these relationships globally generalizable? In this study, we investigated the effects of grazing intensities on ecosystem CO_2_ fluxes by combining a long-term (7-11 years) field experiment conducted in a typical steppe and a meta-analysis of global grasslands. The objectives of this study were to: (i) investigate the effects of grazing intensity with annual wetness fluctuations on ecosystem CO_2_ fluxes (GPP, ER and NEP) covering the 7th to 11th years of a continuous grazing experiment in a typical steppe as well as the meta-analysis in global grasslands; (ii) explore how environmental factors (particularly wetness index, soil moisture and temperature, and grazing intensity) regulate the effects of grazing on ecosystem CO_2_ fluxes.

## 2. Materials and methods

### 2.1. Experimental site

The grazing experiment was conducted at Xilinhot National Climate Observatory (44°12′ N, 116°19′ E, 1129 m above sea level), in the middle part of Inner Mongolian typical steppe in China. The average annual precipitation was 291 mm from 2019 to 2023, with average precipitation in the growing season (May to September) varying from 168 to 325 mm (Fig. S1). The average mean air temperature was 3.9 ℃ (from 2019 to 2023). The soil is classified as a Haplic Calcisol based on the FAO soil classification system or a Calcic-Orthic Aridisol based on the USDA soil classification system (Liang et al., 2021). The vegetation community is dominated by *Stipa grandis* P. Smirn (perennial bunchgrass) *and Leymus chinensis* Trin. Tzvel (perennial rhizomatous grass) (Wu et al., 2024).

### 2.2. Experimental design

The grazing experiment was initiated in 2013 and was based on a randomized design (Fig. S2). There were four treatments with three replicates: control without grazing (CK), light grazing (LG, 0.64 sheep ha^-1^ year^-1^), moderate grazing (MG, 1.28 sheep ha^-1^ year^-1^) and heavy grazing (HG, 2.56 sheep ha^-1^ year^-1^) conducted in plots with 1.44 ha (120 ×120 m^2^) established in the research station (Fig. S2). Two-year-old sheep similar in size were selected to use for grazing from May to September in each year. Sheep were grazing from 7:00 AM to 6:00 PM each day, and then gathered into an enclosure at night (Liang et al., 2021).

### 2.3. Measurement of ecosystem CO_2_ fluxes

Ecosystem CO_2_ fluxes were measured using a transparent chamber (0.5 × 0.5 × 0.8 m^3^) connected to an infrared gas analyzer (LI-8100, Lincoln, NE, USA). To ensure an airtight seal during measurements, the junction between the chamber bottom and the aluminum base was secured with tapes. After positioning the chamber, the enclosed air was continuously mixed by two fans in the chamber. The CO_2_ concentrations were recorded 12 times at 10-second intervals over a 2-minute period. The NEP was then determined from the rate of CO_2_ concentration change, combined with chamber volume and headspace temperature data. A positive NEP value indicated net ecosystem CO_2_ uptake, whereas a negative value denoted net ecosystem CO_2_ release. Following each NEP measurement, the chamber was removed to vent the headspace and then covered with a dark cloth to inhibit photosynthesis before being repositioned on the same base for ER measurement. Both the NEP and ER were measured throughout the growing seasons from 2019 to 2023, corresponding to the 7th to 11th years of grazing. The GPP was thus calculated as the sum of NEP and ER using data collected over this five-year period. The gas exchange measurements were conducted on sunny and calm days between 9:00 and 11:00, a time when the ecosystem CO_2_ fluxes represented the daily average (Niu et al., 2008; Rong et al., 2017; Yu et al., 2025), with a frequency of three times per month. In each plot, the chambers of two replicates were measured. Thus, six replicates per treatment were set up in the field experiment.

### 2.4. Climate conditions and soil microclimate

The annual precipitation (AP) and mean annual air temperature (MAT) for the field study area from 2019 to 2023 were obtained from the China Meteorological Data Service Center (http://data.cma.cn/) of the Xilinhot Station, which was near the experimental site in this field experiment. The De Martonne wetness index (WI) was calculated as follows (De Martonne, 1926; Song et al., 2019):

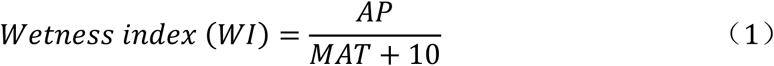

Soil temperature and moisture at a depth of 0-10 cm were measured simultaneously with ecosystem CO_2_ flux measurements, using the probes of the LI-8100 system. Soil moisture was the soil volumetric water content. Measurement sites for soil temperature and moisture were selected near the collars randomly placed in the plots. They were investigated synchronized with the measurements of ecosystem CO_2_ fluxes.

### 2.5. Plant sampling and analyses

Four grazing rotations were conducted each year from 2019 to 2023 in the field study. Before each grazing rotation, two cages (1.2 m × 1.2 m ×1.2 m) were installed in each plot to ensure that the vegetation inside would not be foraged by sheep. Aboveground plant biomass was collected after the end of each grazing rotation. Specifically, in each plot, five 1 m ×1 m quadrats were placed. All AGB within these quadrats was clipped at ground level and then oven-dried at 65 °C for 48 h to determine the community-level dry biomass. The plant species were classified into C_3_ and C_4_ groups. Belowground biomass (BGB) was quantified by collecting root biomass using soil core with diameter of 7 cm each September from 2019 to 2023 (Fig. S2). Specially, two soil cores were collected from each 1 m × 1 m quadrat corresponding to BGB measurements at depths of 0-30 cm. The belowground parts were then extracted from the soil by washing with water and oven-dried at 65 °C for 48 h to obtain dry weight.

The relative growth rate (RGR) was defined as the ratio of aboveground biomass regeneration to the number of days between two consecutive grazing events and was calculated by Eq. (2) (Zong & Shi, 2019):

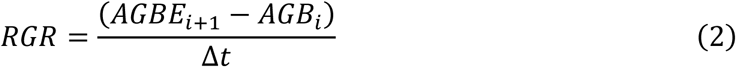

where *AGBE_i+_*_1_: the AGB inside the cages after the (*i*+1)-th grazing round, *AGB_i_*: the remaining AGB after the *i*-th grazing, △*t*: the number of days between the *i*-th and (*i*+1)-th grazing rounds. Aboveground net primary productivity (ANPP) was estimated as the sum of the total herbage consumed by livestock throughout the growing season and the AGB measured at the end of the final grazing rotation of the season.

### 2.6. Data Analysis

All the analyses were conducted using the R (version 4.4.3) (R Core Team, 2023). Annual mean values were calculated for GPP, ER, NEP, AGB, BGB, root/shoot ratio, soil temperature, and soil moisture from 2019 to 2023. Differences across years and among treatments within each year were examined using the Least Significant Difference (LSD) test. Response ratios of GPP, ER, NEP, AGB, BGB, root/shoot ratio, soil temperature, and soil moisture were calculated based on Eq. (3). Additionally, linear regression analysis was conducted using the “*lm*” function to evaluate the relationships of GPP, NEP, AGB, BGB, and RGR with WI.

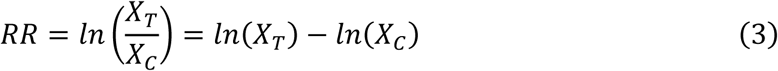

Where *_XT_* and *_XC_* are the mean values of the abovementioned indicators for the grazing (T) and control (C) treatments, respectively.

The repeated measures ANOVA (the *ezANOVA* function of the “*ez*” R package) was used to examine the effects of grazing treatment and sampling years on ecosystem CO_2_ fluxes (GPP, ER, and NEP), microclimatic variables (soil temperature and moisture), and plant biomass parameters (AGB, BGB, and root/shoot ratio) across the five-year study period from 2019 to 2023 (Chen et al., 2024; Lawrence et al., 2026).

We also conducted a random forest model to identify the significant environmental predictors of GPP, ER NEP, AGB and BGB (Archer, 2025; Liaw & Wiener, 2002; Liao et al., 2024). The environmental predictors considered in the field study included grazing intensity, the response ratio of soil temperature and soil moisture, wetness index, and grazing duration. To evaluate the direct and indirect effects of grazing intensity, wetness index, soil temperature, soil moisture, RGR, AGB, and C_3_ and C_4_ plant biomass and richness on NEP, we conducted a structural equation modeling (SEM) analysis using the ‘*piecewiseSEM*’ packages. A good fit of the SEM was evaluated using Fisher’s C value, which was non-significant (*P* > 0.05) (Lefcheck, 2016;Yang et al., 2025).

### 2.7. Meta-analysis of grazing experiments in global grasslands

To assess the effects of grazing on GPP, ER, and NEP in global grasslands, we conducted a systematic literature search for relevant papers published up to June 8, 2025, using the Web of Science (http://apps.webofknowledge.com/), China National Knowledge Infrastructure (CNKI, https://www.cnki.net) and Google Scholar (https://scholar.google.com/). The keywords and terms used were (“grazing” OR “stocking” OR “livestock” OR “grazing intensity”) AND (“net ecosystem productivity” OR “net ecosystem exchange” OR “gross ecosystem productivity” OR “ecosystem respiration” OR “soil carbon flux” OR “soil CO_2_” OR “soil carbon flux” or “soil carbon emission”). Relevant studies were selected based on the following criteria (Fig. S9): (a) Studies were conducted using simulating grazing experiments, and studies with trampling and mowing were excluded. (b) The control and grazing plots were under similar conditions, such as dominant species, community composition, soil type and climate. (c) Data from non-grazed treatments were excluded when additional treatments (e.g., fertilization, experimental warming, or precipitation manipulation) were present. (d) The mean and sample size for any measurements of GPP, ER, and NEP under both grazing and control conditions must be reported. (e) The classifications of grazing intensity (light, moderate, and heavy grazing) were primarily based on the definitions provided in the original studies. For studies in which grazing intensity was not explicitly reported, we classified grazing intensity according to the modified Grazing Intensity Classes proposed by the USDA (https://www.fs.usda.gov/Internet/FSE_DOCUMENTS/stelprdb5109714.pdf) (Yin et al., 2023). Finally, 62 published papers with 585 observations of GPP, ER, and NEP responses (*n* = 161, 249, and 175, respectively) were selected (Fig. 4), and the geographical location of these observations were presented in Fig. 1. Meanwhile, the study site, elevation, mean annual precipitation (MAP), MAT, and grazing intensity from the selected papers were recorded. The wetness indices based on MAP and MAT of each site were calculated according to Eq. (1).

Where available, data for soil temperature, soil moisture, AGB, BGB, and root/shoot ratio were also recorded. In addition, to better explore the effects of grazing intensity and environmental factors on ecosystem CO_2_ fluxes, the data was classified into different subgroups by grazing intensity (LG, MG, and HG), mean annual temperature (≤5 ℃ and >5 ℃), wetness index (≤ 30 and > 30), and precipitation (≤ 400 mm and > 400 mm), grassland type (desert grassland, temperate grassland, alpine grassland, savanna, and others), and livestock type (cattle, sheep, and mix).

The effect size of grazing on each variable was quantified using the natural logarithm of response ratios (*RR*) based on Eq. (3) (Feng et al., 2023). In our dataset, standard deviation or standard error was not reported in 242 of the 585 ecosystem CO_2_ fluxes observations. Like in previous studies, the weighing of each observation (*_WRR_*) was calculated as (Pittelkow et al., 2015):

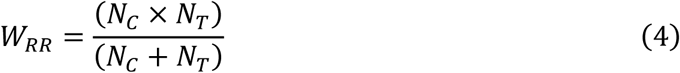

where *_NT_* and *_NC_* are the replicate numbers of treatment and control, respectively.

A linear mixed-effects model, with ‘study’ included as a random factor, was employed to estimate the weighted response ratio (*RR*_++_) across studies or within a specific group, fitting with restricted maximum likelihood using the ‘*lmer*’ function in the ‘lme4’ package (Bates et al., 2015; Feng et al., 2023).

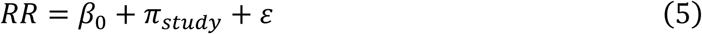

where *_β_*_0_ is the coefficient, *_πstudy_* denotes the random effect associated with ‘study’ (accounting for autocorrelation among observations from the same study), and *ɛ* corresponds to the residual sampling error. We checked the normality of the model residuals using the ‘*check_normality*’ function in the ‘performance’ package. When the assumption of normality was violated, bootstrapping with 999 iterations was performed using the ‘boot’ package to derive the 95% confidence interval (CI) for each *RR_++_* (Chen et al., 2021; Canty et al., 2025). The percentage change (%) of the response ratio was calculated as follows:

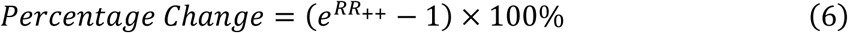

The *lnRR*_++_ was considered statistically significant if its 95% confidence interval (CI) did not overlap zero (Chen et al., 2021).

Potential publication bias for each response variable based on data from the entire database was examined using Egger’s test (Egger et al., 1997). No significant publication bias was detected for any of the variables (Table S4). The random forest model was conducted using “*randomForest*” and “*rfPermute*” packages to identify the relative importance of the predictor variables (wetness index, grazing duration and intensity, the responses of soil moisture and temperature and livestock types) of GPP, ER, NEP, AGB and BGB in the meta-analysis (Archer, 2025; Liaw & Wiener, 2002; Liao et al., 2024). In this model, the importance of each predictor was evaluated based on the percentage increase in the mean squared error (MSE) when that predictor was permuted; a higher MSE value indicated greater importance of the factor. The MSE values for each decision tree, based on out-of-bag estimates from the random forest model, were generated using the “*rfPermute*” package to assess the relative importance of the predictor variables (Liao et al., 2024).

## 3. Results

### 3.1 Grazing intensity effects on ecosystem CO_2_ fluxes and plant biomass in the field experiment

Continuous heavy grazing reduced the mean GPP by 29.87% and the mean ER by 30.94% in the grazing experiment; however, NEP did not differ between HG and CK (Fig. 2). The mean GPP and ER were decreased by 17.46% and 22.72%, respectively, in MG than in CK, but NEP was not different between MG and CK (Table S2). The mean ER was decreased by 21.49% and 29.84%, respectively, in MG and HG than in LG. The mean NEP was decreased by 37.41% in HG than in LG, but was not different among LG, MG and CK. The GPP and NEP were 29.27% and 85.83% higher, respectively, in LG than in CK, but the ER was not different in 2020, the year with the highest precipitation and wetness index among the five years (Fig. S1 and 2). The mean aboveground and belowground biomass was decreased by 39.27% and 19.44%, respectively, in HG than in CK, but the mean root/shoot ratio was increased by 30.59% in HG than in CK (Fig. 2; *P* < 0.05).

**Fig. 2.**
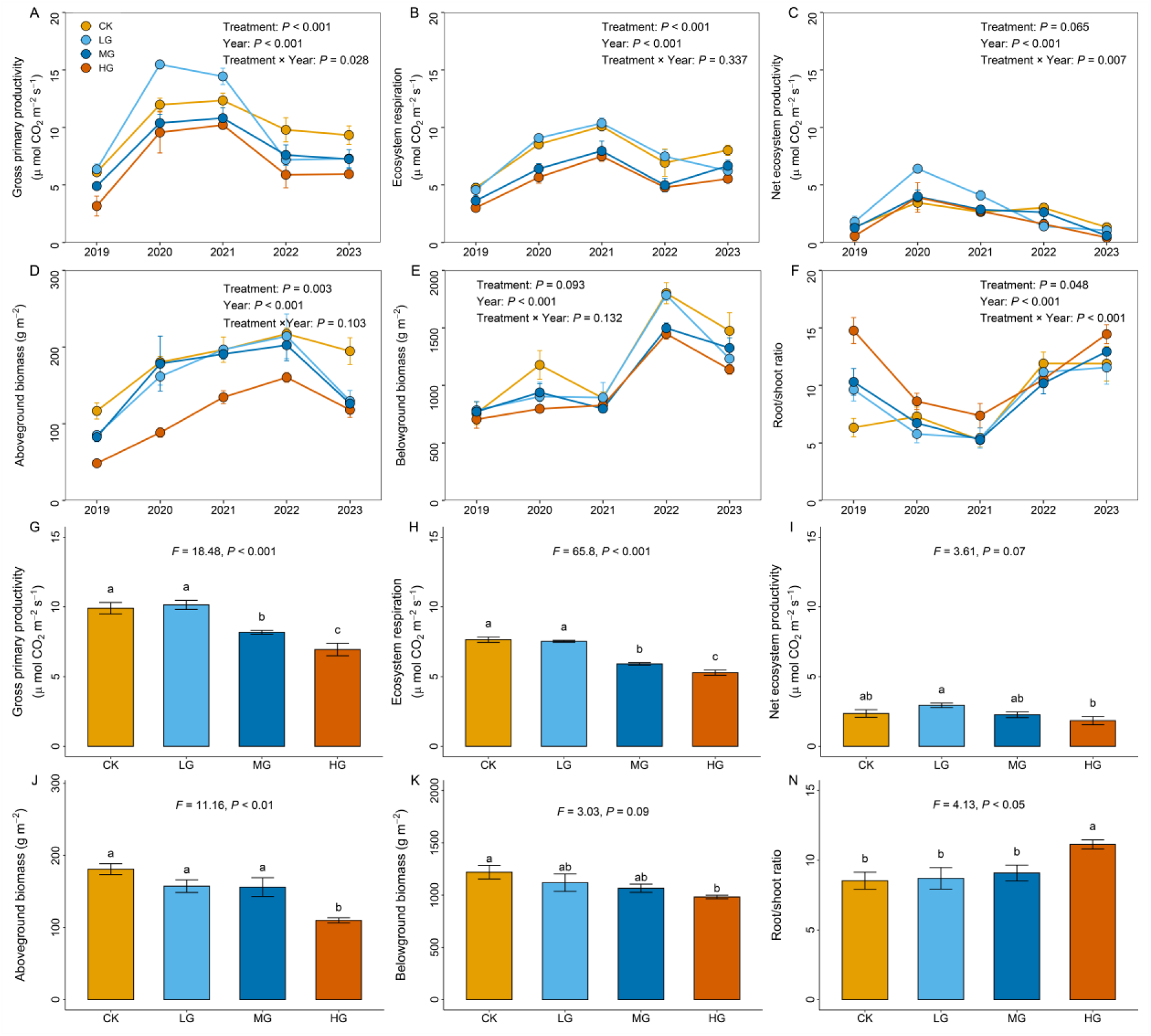
Effects of different grazing intensities on ecosystem carbon dioxide (CO_2_) fluxes and biomass in our field experiment. CK, without grazing; LG, light grazing; MG, moderate grazing; HG, heavy grazing. Data (means ±SE, *n* = 3) followed by different lowercase letters indicate differences at *P* < 0.05 between treatments.

Wetness was the most important predictor of NEP, while grazing intensity was the most important predictor of GPP and ER, revealing a strong coupling between CO_2_-driven changes in wetness index and grazing intensity (Fig. S3). Grazing intensity and duration were also the important predictors influencing AGB and BGB. The linear regression analysis showed that overall grazing as well as light grazing stimulated the responses of GPP and NEP with increasing wetness index (Fig. 3 and Table S3; *P* < 0.05). This relationship could be explained by the significant positive correlation between NEP and both ANPP and RGR (Fig. S4). In the wettest year of 2020, the relative growth rate peaked relative to other years, reaching levels 3.16, 2.76 and 1.96 times higher under LG, MG, and HG respectively, compared to CK (Fig. S5). The NEP increased with WI under HG (Fig. S6; *P* < 0.01), whereas the response ratio of NEP to heavy grazing was not significantly correlated with WI (Fig. 3B). Grazing reduced the response of GPP, ER, AGB, BGB and soil moisture in the field experiment (Fig. 4A). Heavy grazing decreased C_3_ plant biomass but increased C_4_ plant richness compared with other treatments (Fig. S7). In addition, the structural equation model showed that grazing altered net ecosystem productivity by increasing plant relative growth rate, while decreasing aboveground biomass of plants (Fig. S8).

**Fig. 3.**
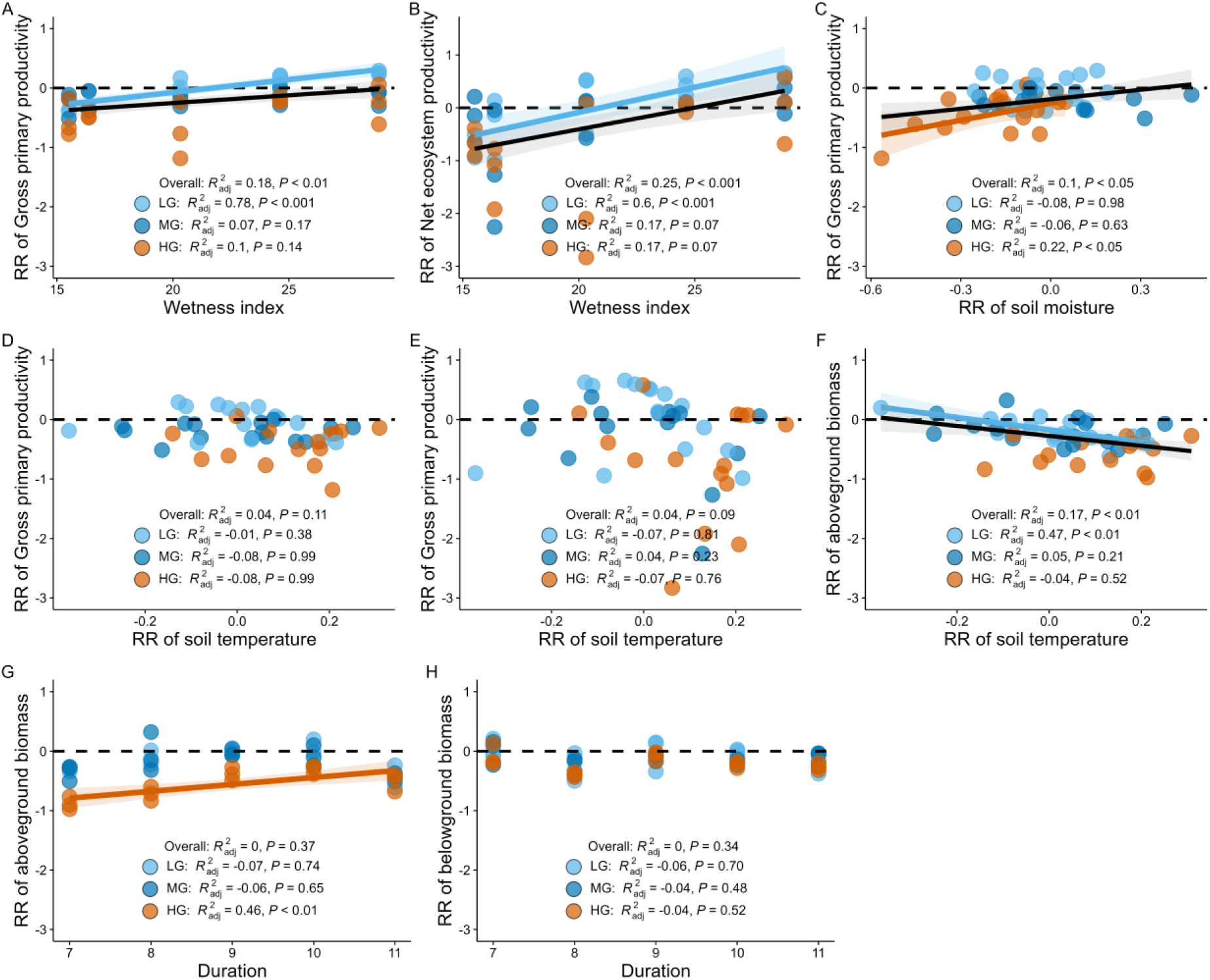
Relationships of the responses of gross primary productivity (GPP) and net ecosystem productivity (NEP) with potential regulating factors from 2019 to 2023 under different grazing intensity. Overall, overall grazing included light, moderate and heavy grazing; LG, light grazing; MG, moderate grazing; HG, heavy grazing. The significant regression lines and 95% confidence intervals are shown with lines (solid for significant) and shaded areas, respectively.

**Fig. 4.**
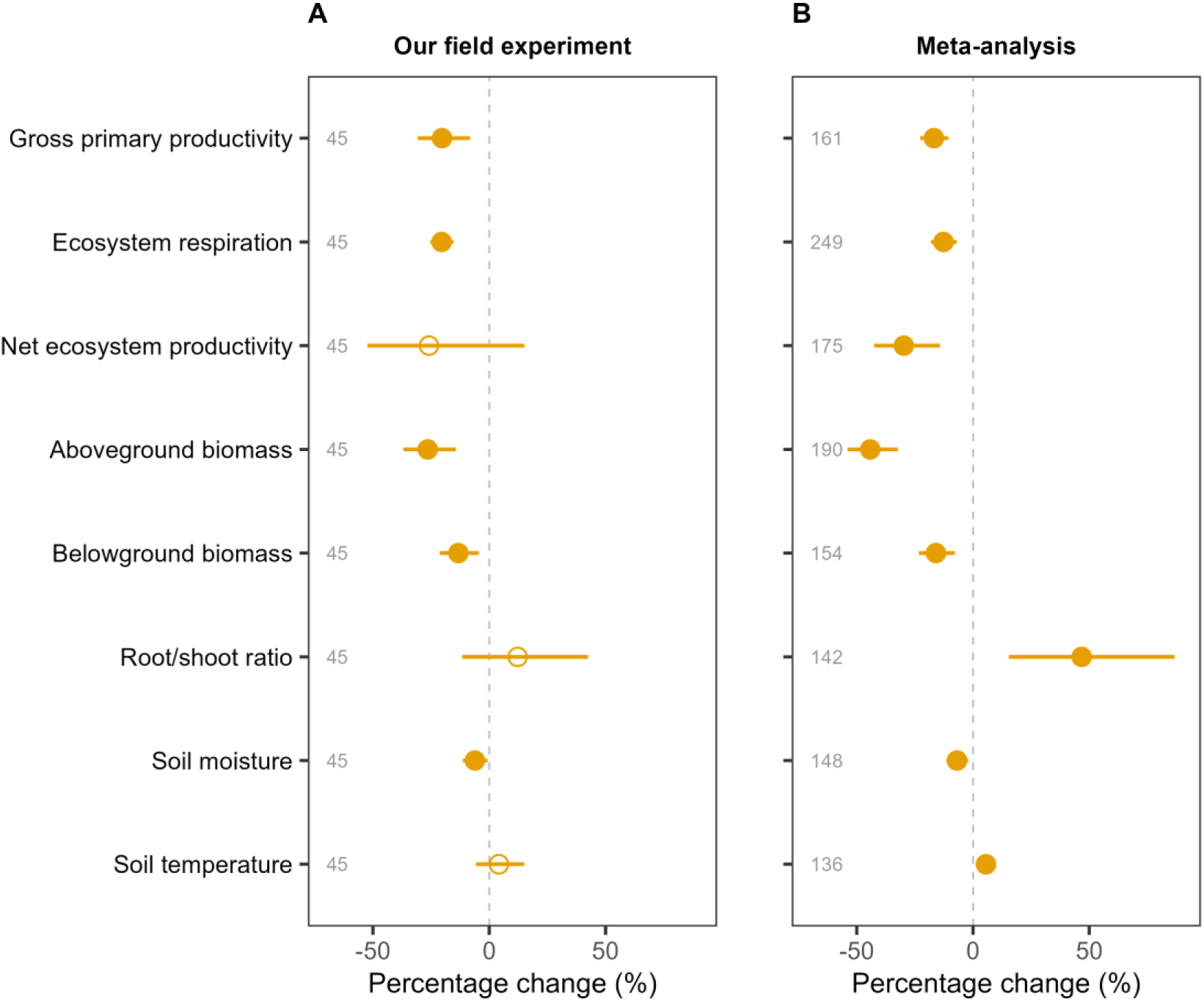
Effects of grazing on gross primary productivity, ecosystem respiration, net ecosystem productivity, aboveground biomass, belowground biomass and root/shoot ratio, soil moisture and soil temperature in the field experiment of this study (A) and in global grasslands of the meta-analysis (B). Circles and error bars represent average parameter estimates and 95% confidence interval (CI). The solid circle indicates significance when the CI did not overlap zero. Sample sizes of observations for each variable are displayed on the left.

### 3.2 Synthesis of grazing on ecosystem CO_2_ fluxes and plant biomass across global grasslands

In the meta-analysis across global grasslands, grazing significantly decreased ecosystem CO_2_ fluxes by 13.56% to 30.46% (Fig. 4B). Heavy grazing had a more pronounced negative effect on ecosystem CO_2_ fluxes compared to light and moderate grazing (Fig. 5A). The response of GPP was decreased by 15.89%, 11.88% and 26.34%, respectively, under LG, MG and HG. The response of ER was decreased by 7.42%, 14.91% and 21.82%, respectively, under LG, MG and HG. The response of NEP was decreased by 18.86% and 34.89%, respectively, in LG and HG, but was not significant in MG. Light, moderate and heavy grazing reduced AGB by 27.89%, 34.57% and 66.17% (Fig. 5B). Moderate and heavy grazing decreased the BGB by 14.85% and 26.13%, respectively, but the response of BGB was not significant under light grazing. Grazing increased the root/shoot ratio by 31.78% to 111.11%. Heavy grazing had a more pronounced positive effect on root/shoot ratio compared to light and moderate grazing.

**Fig. 5.**
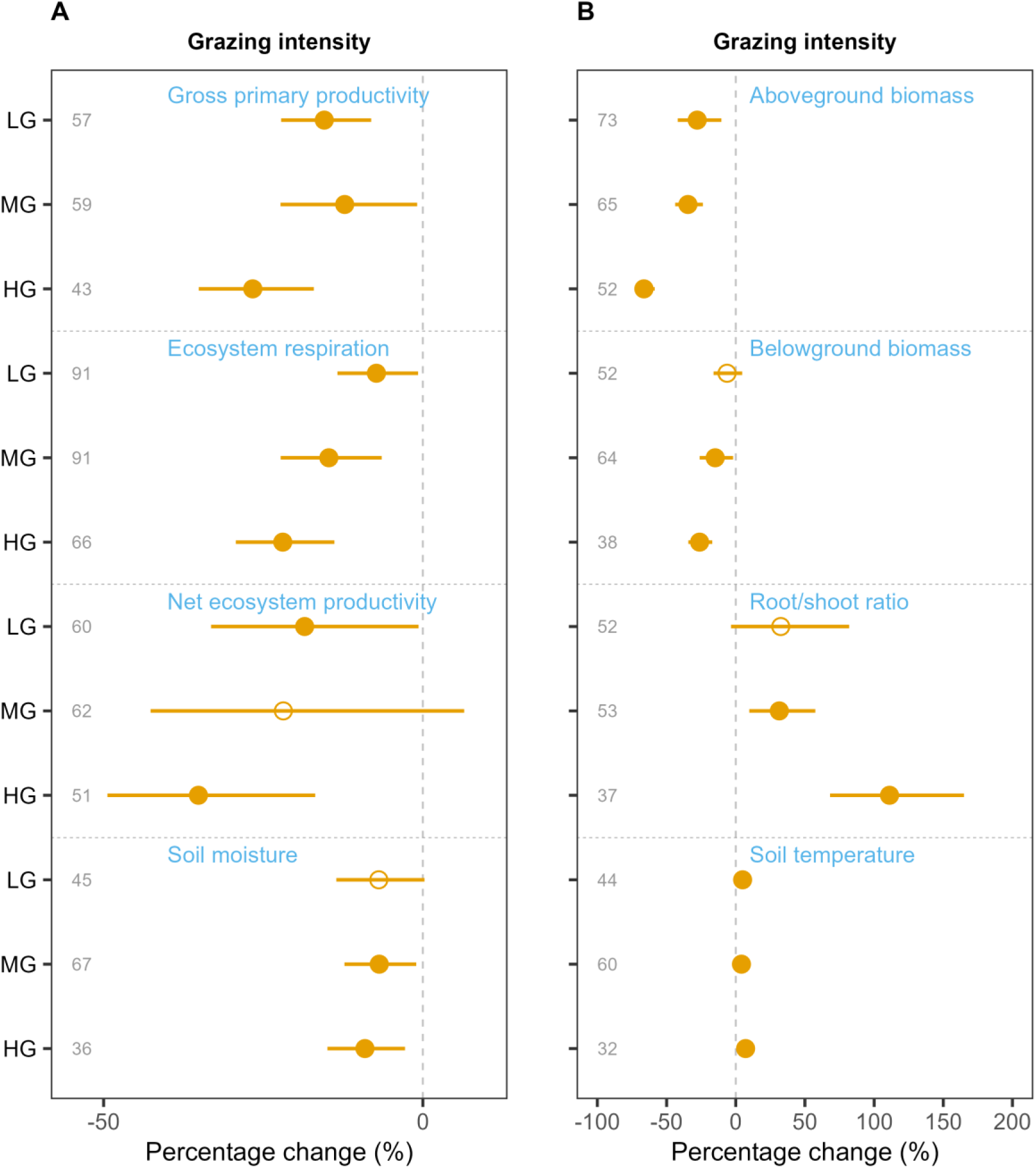
Effects of different grazing intensities on ecosystem CO_2_ fluxes, biomass and soil temperature and moisture in global grasslands from the meta-analysis. Circles and error bars represent average effects and their confidence interval (CI). The solid circle indicates significant effects, i.e., the 95% CI did not overlap zero. The sample size for each variable is shown on the left. LG, light grazing; MG, moderate grazing; HG, heavy grazing.

Globally, grazing decreased GPP, ER and NEP under lower precipitation conditions (≤ 400 mm), while the reductions were not significant under higher precipitation conditions (> 400 mm) (Fig. S10A). Grazing significantly reduced GPP, ER, and NEP at mean annual temperatures ≤ 5 ℃, but these effects were not significant at temperatures > 5 ℃ (Fig. S10B). Similarly, under lower wetness conditions (WI ≤ 30), grazing decreased GPP, ER, and NEP more than under higher wetness conditions (WI > 30) (Fig. 6). The response of NEP was decreased under lower WI (WI ≤ 30) in MG and HG, but it was not different under higher WI (WI > 30) in MG. The response of NEP was decreased under higher WI (WI > 30) in HG. Higher wetness index mitigated the response of NEP to grazing. The duration of grazing also influenced its effect: GPP and ER declined more substantially under longer grazing durations (Fig. S10C). The most pronounced decline in NEP occurred under grazing durations of 5 to 10 years (*P* < 0.001). Grazing decreased GPP, ER and NEP in desert and temperate grassland, but had no significant effect on GPP, ER and NEP in alpine grassland and savanna (Fig. S11). In addition, cattle and sheep grazing decreased GPP and NEP, but livestock mixed grazing did not show significant effect on GPP, ER and NEP (Fig. S12).

**Fig. 6.**
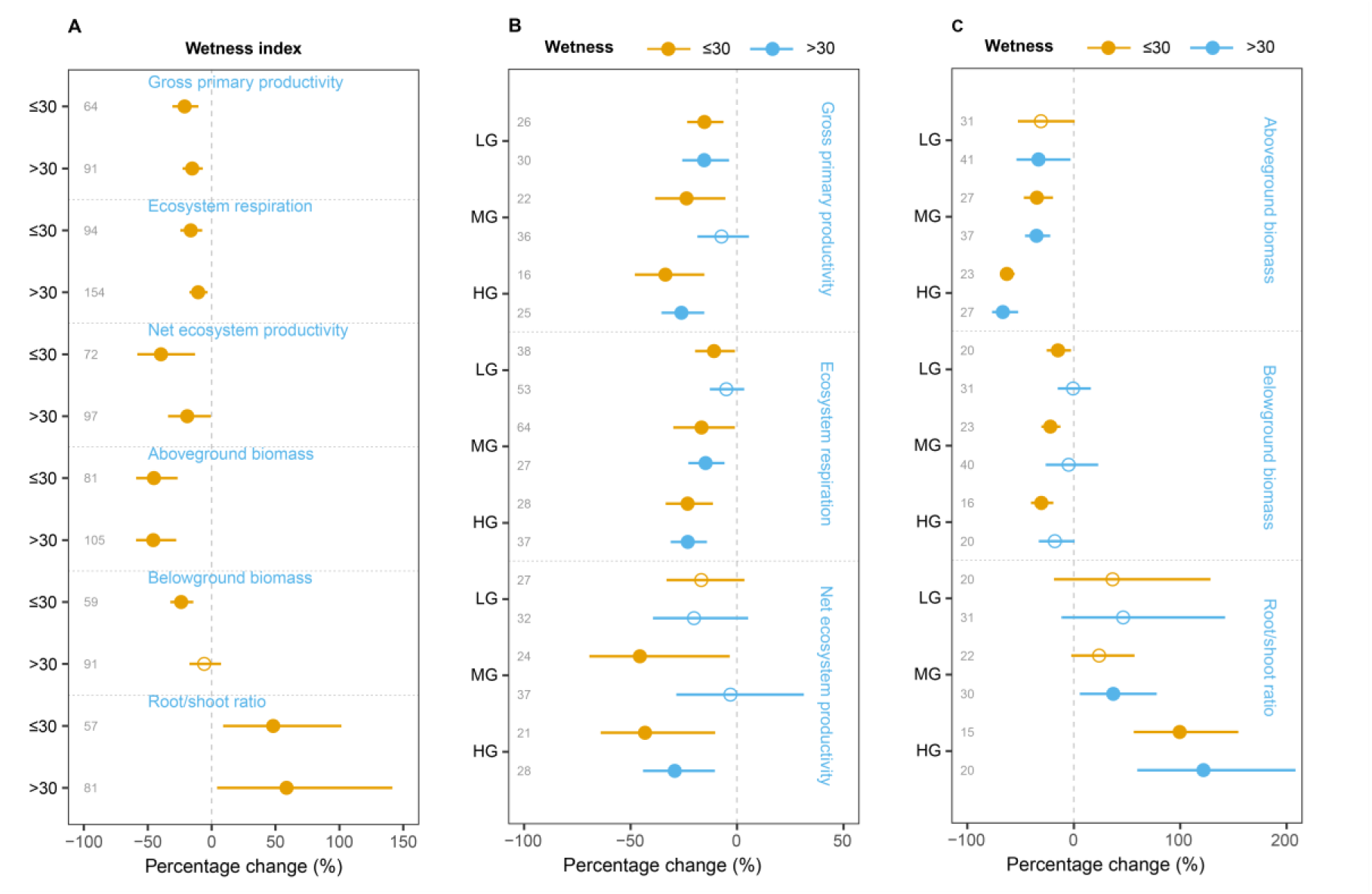
Responses of ecosystem CO_2_ fluxes and biomass to grazing (A) and different grazing intensities (B-C), at wetness index ≤ 30 and wetness index > 30 in global grasslands from the meta-analysis. Circles and error bars represent average parameter estimates and 95% confidence interval (CI). The solid circle indicates significance when the CI did not overlap zero. The sample size for each variable is shown on the left. LG, light grazing; MG, moderate grazing; HG, heavy grazing.

In global grasslands, the key environmental factors influencing GPP and ER response to grazing included WI, grazing duration, the response of soil moisture, and grazing intensity (Fig. S13; *P* < 0.05). For NEP, the major predictors were WI, grazing duration and intensity (*P* < 0.05). The most important environmental parameters for AGB response to grazing were grazing intensity, WI, grazing duration, the response of soil moisture and temperature and livestock types. For BGB, the major predictors were WI, grazing duration, the response of soil moisture and temperature. The response of NEP was positively correlated with WI under light, moderate and total grazing in global grasslands (Fig. 7; *P* < 0.1). The response of ecosystem CO_2_ fluxes was affected by AGB under grazing in global grasslands (Fig. S14).

**Fig. 7.**
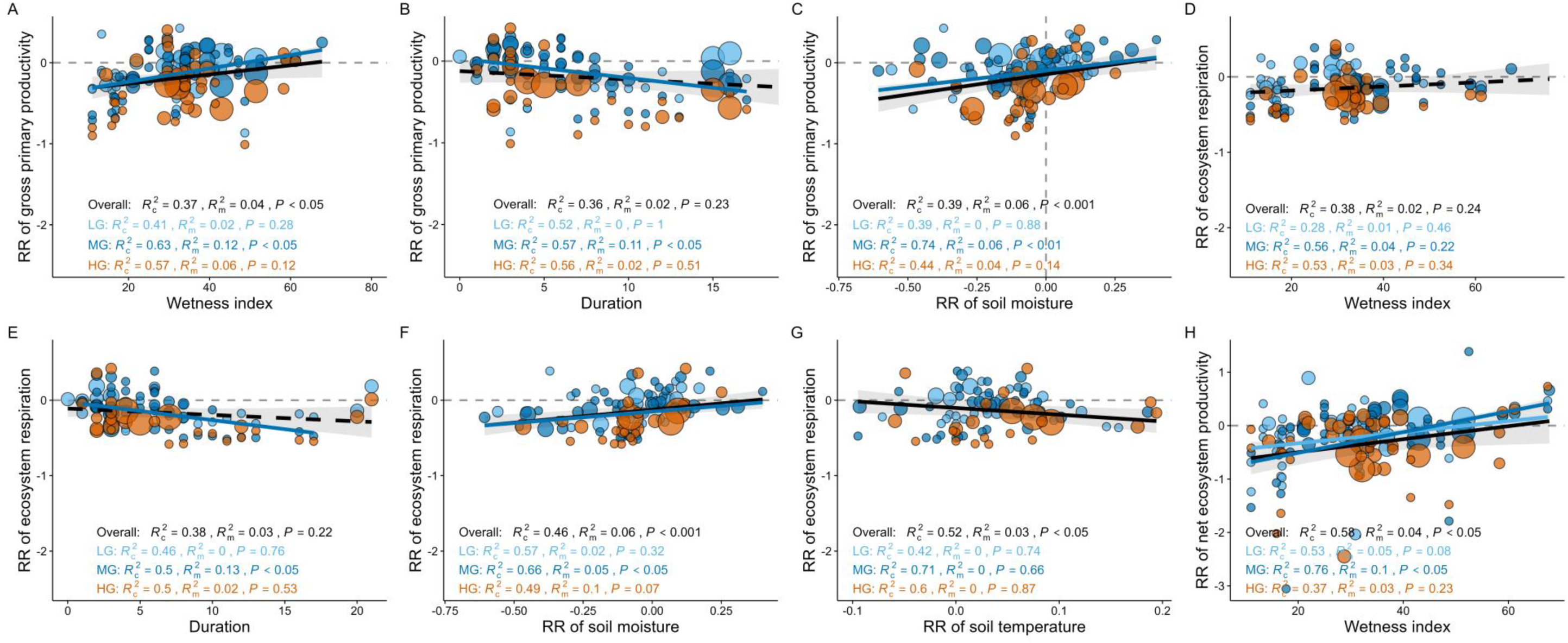
Relationships of the responses of ecosystem CO_2_ fluxes with potential regulating factors in global grasslands from the meta-analysis. The marginal (m) and conditional (c) *R*^2^ indicate the proportion of variance explained by the fixed effects and by both the fixed and random effects (study), respectively. The black solid lines and gray shaded areas in panels are the mean and 95% confidence interval of the slope, respectively. The sizes of points are proportional to their corresponding weights. RR, response ratio; overall, total grazing; LG, light grazing; MG, moderate grazing; HG, heavy grazing.

## 4. Discussion

### 4.1 The responses of ecosystem CO_2_ fluxes and plant biomass to grazing and controlling factors

Grazing decreased the responses of ecosystem CO_2_ fluxes and plant biomass in global grasslands, but only NEP was not significantly affected by grazing in our field experiment (Fig. 4). The lack of a significant response in NEP may be because the site in this field study was managed for year-round continuous low-intensity grazing (Liang et al., 2021). In addition, grazing affected GPP and ER at similar magnitudes, resulting in limited net changes in NEP. Similarly, the reason why the non-significant response of NEP appeared only under MG may be that moderate grazing affected GPP and ER at similar magnitudes (Fig. 5). However, the reduction in response of GPP was higher than that of ER in LG and HG, resulting in the decrease in NEP. Both the results of field experiment and global meta-analysis showed that light grazing did not reduce NEP under wetter conditions (Fig. 3B, 6B and 7H). Light grazing usually stimulates leaf regrowth following defoliation, and these new leaves often are more physiologically active than the older leaves that contribute much of leaf area in ungrazed treatment (Polley et al., 2008), which likely imply a stronger leaf photosynthesis and C sink (Reich et al., 2007). Considering factors such as different grassland types (Fig. S11), livestock grazing modes (Fig. S12), climatic conditions, and grazing durations, the future grazing studies on NEP should focus more on ANPP and RGR, and extrapolate the results cautiously.

Grazing stimulated the NEP under wet conditions, but depressed it under dry conditions, a pattern consistent across the typical steppe in our experiment and supported by our global meta-analysis (Fig. 3B and 7H). This finding aligned with the finding that the response of NEP increased with the increase in wetness index under grazing. This relationship could be explained by the significant positive correlation of NEP with AGB (Fig. S14). The linear regression analysis also showed that light grazing stimulated the responses of NEP with increasing wetness index. This relationship could be explained by the significant positive correlation between NEP and both ANPP and RGR (Fig. S4). Under lower wetness conditions (WI ≤ 30), light grazing reduced GPP and ER to a lesser extent than moderate and heavy grazing (Fig. 6B). Compared with higher grazing intensities, light grazing consumed less forage by livestock, thereby better mitigated the effect of lower moisture on GPP and ER. Moderate grazing decreased GPP and NEP under lower WI, but the responses of GPP and NEP to moderate grazing were not different under higher WI in global grasslands (Fig. 6 and 8), indicating that higher wetness offset the response of GPP and NEP to moderate grazing. This was likely because grazing-induced biomass reduction was less under wet condition in grasslands (Jin et al., 2023). However, not all grazing intensities aligned with the finding that the response of NEP increased with the increase in wetness index, likely due to severe loss of vegetation resilience under heavy grazing. Heavy grazing, even under higher wetness, reduced NEP and AGB in global grasslands (Fig. 5 and 6), resulting in a severe decline or even loss of their resilience, which could not be restored to control levels in long-term heavy grazing (Irob et al., 2023).

**Fig. 8.**
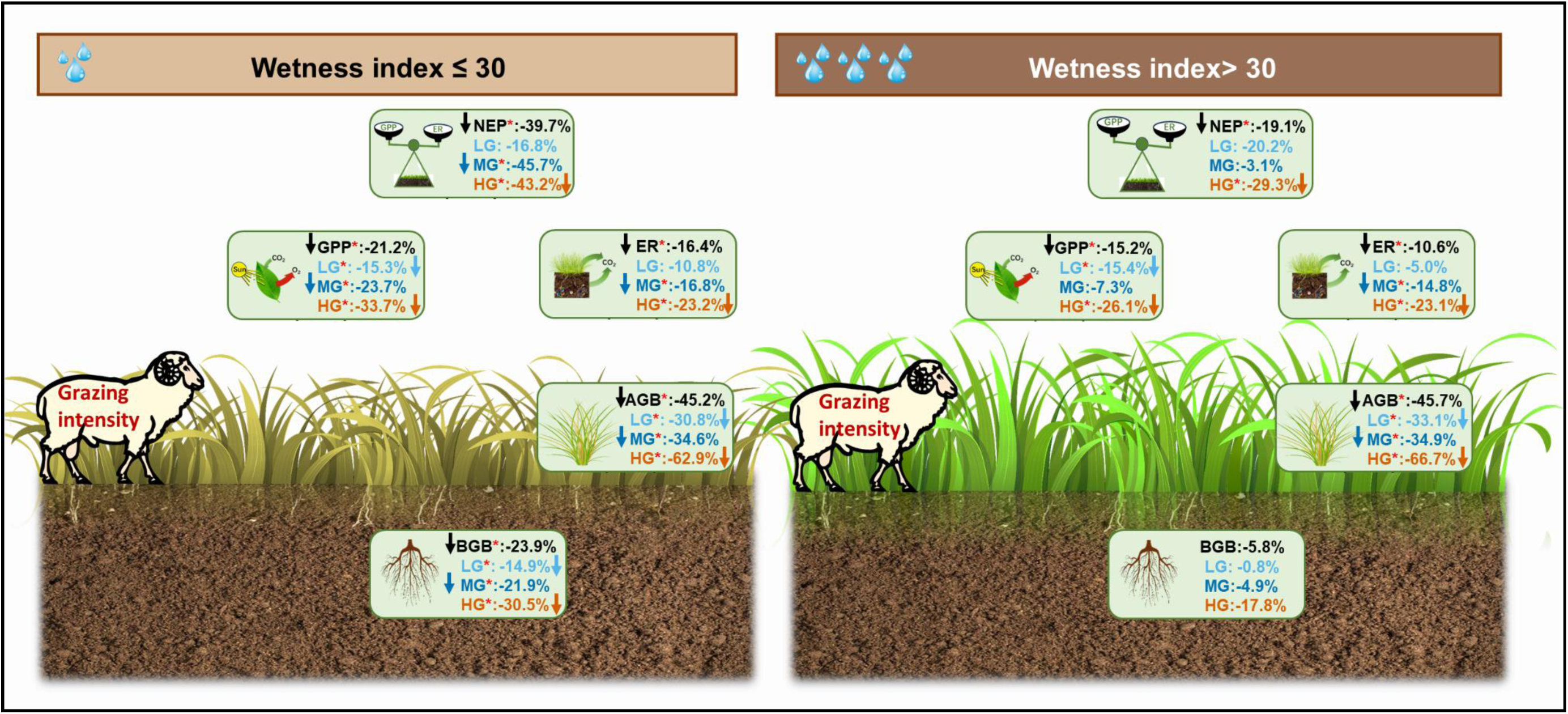
Schematic summary of the effects of grazing intensity on ecosystem carbon dioxide (CO_2_) fluxes and biomass in global grasslands in the meta-analysis. The black and blue arrows indicate negative effects on overall grazing and grazing intensities, respectively. Asterisks (*) indicate significant effects on variables at *P* < 0.05. GPP, gross primary productivity; ER, ecosystem respiration; NEP, net ecosystem productivity; LG, light grazing; MG, moderate grazing; HG, heavy grazing.

Both our experiment and meta-analysis showed that aboveground biomass, grazing intensity and wetness index were key factors regulating the responses of ecosystem CO_2_ fluxes to grazing (Fig. S3, S13 and S14), with their responses intensifying with higher aboveground biomass and wetness index. Although grazing reduced C_3_ plant biomass, it had no significant effects on C_4_ plant biomass or the C_3_:C_4_ plant biomass ratio (Fig. S7). Furthermore, variation in plant community composition did not significantly explain the response of NEP to grazing (Fig. S8), indicating that the photosynthetic pathway composition of the plant community was not the primary driver mediating grazing effects on ecosystem carbon sequestration (Delgado- Balbuena et al., 2013). Grazing reduced aboveground biomass and plant respiration (Liu et al., 2020; Wang et al., 2025), thereby decreasing GPP and ER in both field experiment and meta- analysis (Fig. 4). Plants enhanced photosynthetic structures and physiological functions under grazing pressure. These adaptations improved CO_2_ assimilation and allocated more carbohydrates to compensate for the loss of aboveground biomass, thereby achieving compensatory growth (Wang et al., 2020; Yu et al., 2025). The increase in GPP and NEP in LG in 2020 was due to the increased relative growth rate in LG than in CK (Fig. S5), suggesting that grazing stimulated greater plant compensatory growth in years with more precipitation (Li et al., 2025). Moreover, appropriate grazing intensity facilitated the removal of senescent plant tissues, leading to increased leaf light capture capability and enhanced photosynthetic capacity (Shen et al., 2019). The aboveground and belowground biomass, as well as plant root/shoot ratios were influenced by grazing activities such as feeding, trampling, and excretion (Cao et al., 2024). In the field experiment and global meta-analysis of this study, plant biomass (both aboveground and belowground biomass) was lower under heavy grazing (Fig. 2 and 5). Under heavy grazing conditions, where soil nutrients were often depleted, plants tended to prioritize biomass allocation to roots to enhance nutrient acquisition and utilization (Liu et al., 2024). Moreover, heavy grazing can stimulate C allocation to roots (Ren et al., 2024), which explains why the root/shoot ratio was higher under heavy grazing (Fig. 2 and 5). In the wetter years (2020 and 2021), the aboveground biomass was higher while the root/shoot ratio was lower than in the drier years. Heavy grazing affected the allocation strategy of aboveground and belowground biomass by plants.

Our long-term experiment and global meta-analysis jointly provided complementary evidence for understanding the effect of grazing on the ecosystem CO_2_ fluxes in grassland ecosystems. Long-term grazing field experiments revealed the mechanisms underlying the effects of grazing on ecosystem CO_2_ fluxes. Global meta-analysis further clarified the universality of these response patterns across different climatic zones and grassland types. It should be noticed that both field experiment and meta-analysis consistently revealed that wetness was an important factor in regulating the ecosystem CO_2_ fluxes in response to grazing. Under wetter conditions, light grazing promoted compensatory plant growth, enhanced leaf turnover and photosynthetic recovery capabilities, thereby maintaining or even enhancing the NEP (Morgan et al., 2016; Owensby et al., 2006). Conversely, under drier conditions or heavy grazing pressure, a decrease in aboveground biomass and weakened ecosystem resilience jointly limited the ecosystem carbon uptake capacity. These results suggested that wetness modulated the effects of grazing on NEP in grasslands. Moderate grazing may be sustainable under better water conditions, while heavy grazing may exceed the ecosystem resilience threshold even under relatively wet conditions, leading to sustained ecological degradation.

### 4.2 Limitations and implications for future study

Our results highlight that light grazing serves as a promising management strategy to promote ANPP and CO_2_ sequestration. However, some aspects still need to be further explored. The assessment of ecosystem CO_2_ fluxes was incomplete due to limited measurement time, area and sampling intervals. This may bias the representation of seasonal cycles and spatial heterogeneity, especially in the regions that were not monitored. Multiple time-point sampling should be adopted; the sampling frequency and points should also be increased in the future field study. Furthermore, key intermediate processes, such as soil nutrient availability, changes in community composition, and leaf photosynthetic physiological parameters, should also be investigated in the future. Our meta-analysis has limitations due to the relatively small sample size, which stems from the scarcity of studies across global grasslands exploring the effects of grazing intensity on ecosystem CO_2_ fluxes. The data of ecosystem CO_2_ fluxes from large-scale eddy covariance flux towers were not included in this meta-analysis, because there is an absence of a robust model to convert data between flux tower observations and chamber-based methods. In addition, future studies could conduct more experiments of ecosystem CO_2_ fluxes in South America, Africa and Oceania, which have large amounts of grasslands for grazing, but were rarely investigated. Furthermore, more drought or wetness indices should be investigated in future studies of grazing effects on ecosystem CO_2_ fluxes.

There are also limitations in evaluating the effects of grazing on grassland-livestock ecosystem CO_2_ fluxes in this study. Specifically, carbon emissions derived from animal respiration were not included in this study. Therefore, the results of the field experiment and meta-analysis in this study should not be interpreted as a complete carbon budget assessment of the grassland-livestock ecosystem. Our primary objective was to evaluate the carbon exchange processes between vegetation and soil under grazing disturbance. Thus, the grazing effects reported in the field experiment and meta-analysis of this study were responses at the vegetation-soil ecosystem scale rather than the net carbon balance of the entire grazing system. In previous studies, meta-analysis literatures of effects of grazing intensities on NEP were limited and not adequate. Regardless of these limitations, the results of our field experiment and meta-analysis have significant implications for future studies and grassland production. The primary factors regulating ecosystem CO_2_ fluxes responses to grazing were identified as aboveground biomass, wetness index and grazing intensity. The NEP was positively affected by wetness index in LG, MG and HG. However, in both the typical steppe grazing experiment and the synthesis of global grasslands, the NEP resilience increased with higher wetness index specifically under light grazing, but not under heavy grazing. During wetter years, the NEP was even higher under LG compared to CK, whereas in drier years, no significant difference was observed between LG and CK (Fig. 2). This pattern aligns with the greater RGR and ANPP observed under elevated wetness index in light grazing treatment (Fig. S4). Globally, higher wetness index and aboveground biomass enhanced the response of NEP to grazing (Fig. 3, 7 and S14). However, even under higher wetness conditions, heavy grazing reduced NEP and AGB in global grasslands (Fig. 6), reflecting a severe decline, or even loss of resilience under long-term heavy grazing pressure. These results indicate that the negative impacts of long-term heavy grazing on NEP could not be offset by higher annual wetness index in global grasslands, underscoring the necessity of implementing strict grazing management measures. Additionally, moderate grazing under wetter years may be an appropriate management strategy to sustain grassland carbon balance while supporting livestock production.

## 5. Conclusion

In summary, the meta-analysis of this study presents the first comprehensive assessment of how annual wetness index affects the response of ecosystem CO_2_ fluxes to grazing across global grasslands. Our results showed that wetness index and aboveground biomass were key factors regulating the responses of ecosystem CO_2_ fluxes, and both exhibiting positive associations with these responses. Grazing reduced the response of GPP and ER in our field experiment of typical steppe and in the meta-analysis of global grasslands. Higher wetness indices offset the effect of moderate grazing rather than heavy grazing on NEP in global grasslands. These findings underscore the importance of developing region-specific grazing strategies guided by local wetness indices.

## Acknowledgements

The authors would like to acknowledge funding from the National Natural Science Foundation of China (U22A20559, 32260289 and 32271656), the Inner Mongolian Key Research and Development and Achievement Transformation Plan Project (2025YFDZ0062), the Science and Technology Innovation Major Demonstration Project of Inner Mongolia (2024JBGS0007), the Natural Science Foundation of Inner Mongolia (2025MS03013 and 2025MS03119), the State Key Laboratory of Vegetation Structure, Function and Construction (VegLab2025002), and the Fundamental Research Funds for the Directly-Affiliated Universities of the Autonomous Region (2026JBKY044). We would also like to acknowledge the staff in the National Climate Observatory in Xilinhot of Inner Mongolia, China for their help in data collection.

## References

Archer E. (2025) *rfPermute*: Estimate Permutation p-Values for Random Forest Importance Metrics. *R package version 2.5.5* https://CRAN.R-project.org/package=rfPermute

Bardgett, R. D., & Wardle, D. A. (2003) Herbivore-mediated linkages between aboveground and belowground communities. Ecology 84: 2258–2268 10.1890/02-0274

Bates, D., Mächler, M., Bolker, B., & Walker, S. (2015) Fitting Linear Mixed-Effects Models Using lme4. J Stat Softw 67: 1–48 10.18637/jss.v067.i01

Canty A, Ripley B. (2025) *boot*: Bootstrap Functions. R package version 1.3-32 https://CRAN.R-project.org/package=boot.

Cao, F. F., Li, W. B., Jiang, Y., Gan, X. L., Zhao, C. Y., & Ma, J. C. (2024) Effects of grazing on grassland biomass and biodiversity: A global synthesis. Field Crop Res 306: 108852 10.1016/j.fcr.2023.109204

Chang, Q., Xu, T. T., Ding, S. W., Wang, L., Liu, J. S., Wang, D. L., Wang, Y., Li, Z. Q., Zhao, X., Song, X. X., & Pan, D. F. (2020) Herbivore Assemblage as an Important Factor Modulating Grazing Effects on Ecosystem Carbon Fluxes in a Meadow Steppe in Northeast China. J Geophys Res-Biogeosci 125: e2020JG005652 10.1029/2020jg005652

Chen, W. N., Wang, S., Wang, J. S., Xia, J. Y., Luo, Y. Q., Yu, G. R., & Niu, S. L. (2023) Evidence for widespread thermal optimality of ecosystem respiration. Nat Ecol Evol 7: 1379–1387 10.1038/s41559-023-02121-w

Chen, X. L., Chen, H. Y. H., Searle, E. B., Chen, C., & Reich, P. B. (2021) Negative to positive shifts in diversity effects on soil nitrogen over time. Nat Sustain 4: 225–232 10.1038/s41893-020-00641-y

Chen, Y., Qin, W. K., Zhang, Q. F., Wang, X. D., Feng, J. G., Han, M. G., Hou, Y. H., Zhao, H. Y., Zhang, Z. H., He, J. S., Torn, M. S., & Zhu, B. (2024) Whole-soil warming leads to substantial soil carbon emission in an alpine grassland. Nat Commun 15: 4489 10.1038/s41467-024-48736-w

De Martonne, E. (1926) Une nouvelle fonction climatologique: L’indice d’aridité. La Météorologie 2: 449–458

Delgado-Balbuena, J., Arredondo, J. T., Loescher, H. W., Huber-Sannwald, E., Chavez-Aguilar, G., Luna- Luna, M., & Barretero-Hernandez, R. (2013) Differences in plant cover and species composition of semiarid grassland communities of central Mexico and its effects on net ecosystem exchange. Biogeosciences 10: 4673–4690 10.5194/bg-10-4673-2013

Du, C. J., Zhou, G. Y., & Gao, Y. H. (2022) Grazing exclusion alters carbon flux of alpine meadow in the Tibetan Plateau. Agric For Meteorol 314: 108774 10.1016/j.agrformet.2021.108774

Egger, M., Davey Smith, G., Schneider, M., & Minder, C. (1997) Bias in meta-analysis detected by a simple, graphical test. BMJ 315: 629–634 10.1136/bmj.315.7109.629

Feng, J. G., Song, Y. J., & Zhu, B. (2023) Ecosystem-dependent responses of soil carbon storage to phosphorus enrichment. New Phytol 238: 2363–2374 10.1111/nph.18907

Lawrence, M. A. (2026) ez: Easy Analysis and Visualization of Factorial Experiments. R package version 4.5–0 https://CRAN.R-project.org/package=ez

Lefcheck, J.S. (2016) piecewiseSEM: Piecewise structural equation modelling in r for ecology, evolution, and systematics. Methods Ecol Evol 7: 573–579 10.1111/2041-210X.12512

Liaw, A., & Wiener, M. (2002) Classification and regression by randomForest. R news 2: 18–22 https://CRAN.R-project.org/doc/Rnews/

Lüdecke, D., Ben-Shachar, M. S., Patil, I., Waggoner, P., & Makowski, D. (2021) performance: An R Package for Assessment, Comparison and Testing of Statistical Models. J Open Source Softw 6: 3139 10.21105/joss.03139

Gomez-Casanovas, N., DeLucia, N. J., Bernacchi, C. J., Boughton, E. H., Sparks, J. P., Chamberlain, S. D., & DeLucia, E. H. (2018) Grazing alters net ecosystem C fluxes and the global warming potential of a subtropical pasture. Ecol Appl 28: 557–572 10.1002/eap.1670

Hao, Y. B., Zhang, H., Biederman, J. A., Li, L. F., Cui, X. Y., Xue, K., Du, J. Q., & Wang, Y. F. (2018) Seasonal timing regulates extreme drought impacts on CO_2_ and H_2_O exchanges over semiarid steppes in Inner Mongolia, China. Agric Ecosyst Environ 266: 153–166 10.1016/j.agee.2018.06.010

He, M., Zhou, G. Y., Yuan, T. F., van Groenigen, K. J., Shao, J. J., & Zhou, X. H. (2020) Grazing intensity significantly changes the C : N : P stoichiometry in grassland ecosystems. Glob Ecol Biogeogr 29: 355–369 10.1111/geb.13028

Heimann, M., & Reichstein, M. (2008) Terrestrial ecosystem carbon dynamics and climate feedbacks. Nature 451: 289–292 10.1038/nature06591

Hilker, T., Natsagdorj, E., Waring, R. H., Lyapustin, A., & Wang, Y. J. (2014) Satellite observed widespread decline in Mongolian grasslands largely due to overgrazing. Glob Change Biol 20: 418–428 10.1111/gcb.12365

Hou, L. Y., Liu, Y., Du, J. C., Wang, M. Y., Wang, H., & Mao, P. S. (2016) Grazing effects on ecosystem CO_2_ fluxes differ among temperate steppe types in Eurasia. Sci Rep 6: 29028 10.1038/srep29028

Irob, K., Blaum, N., Weiss-Aparicio, A., Hauptfleisch, M., Hering, R., Uiseb, K., & Tietjen, B. (2023) Savanna resilience to droughts increases with the proportion of browsing wild herbivores and plant functional diversity. J Appl Ecol 60: 251–262 10.1111/1365-2664.14351

Jiang, Z. Y., Hu, Z. M., Lai, D. Y. F., Han, D. R., Wang, M., Liu, M., Zhang, M., & Guo, M. Y. (2020) Light grazing facilitates carbon accumulation in subsoil in Chinese grasslands: A meta-analysis. Glob Change Biol 26: 7186–7197 10.1111/gcb.15326

Jin, Y. X., Tian, D. S., Li, J. W., Wu, Q., Pan, Z. L., Han, M. Q., Wang, Y. H., Zhang, J., & Han, G. D. (2023) Water causes divergent responses of specific carbon sink to long-term grazing in a desert grassland. Sci Total Environ 873: 162166 10.1016/j.scitotenv.2023.162166

Ju, X., Wang, B., Wu, L., Zhang, X., Wu, Q., & Han, G. (2024) Grazing decreases net ecosystem carbon exchange by decreasing shrub and semi-shrub biomass in a desert steppe. Ecol Evol 14: e11528 10.1002/ece3.11528

Knauer, J., Cuntz, M., Smith, B., Canadell, J. G., Medlyn, B. E., Bennett, A. C., Caldararu, S., & Haverd, V. (2023) Higher global gross primary productivity under future climate with more advanced representations of photosynthesis. Sci Adv 9: eadh9444 10.1126/sciadv.adh9444

Lemoine, N. P., Sheffield, J., Dukes, J. S., Knapp, A. K., & Smith, M. D. (2016) Terrestrial Precipitation Analysis (TPA): A resource for characterizing long-term precipitation regimes and extremes. Methods Ecol Evol 7: 1396–1401 10.1111/2041-210x.12582

Li, Y. L., Li, F. Y., Shi, C. J., Wang, H., Wu, L., Wang, Y. D., & Minggagud, H. (2025) Precipitation and grazing intensity jointly shape plant compensatory growth and productivity in a semi-arid steppe ecosystem. Agric Ecosyst Environ 393: 109834 10.1016/j.agee.2025.109834

Liang, M. W., Smith, N. G., Chen, J. Q., Wu, Y. T., Guo, Z. W., Gornish, E. S., & Liang, C. Z. (2021) Shifts in plant composition mediate grazing effects on carbon cycling in grasslands. J Appl Ecol 58: 518–527 10.1111/1365-2664.13824

Liao, H. P., Liu, C., Zhou, S. G., Liu, C. Q., Eldridge, D. J., Ai, C. F., Wilhelm, S. W., Singh, B. K., Liang, X. L., Radosevich, M., Yang, Q. E., Tang, X., Wei, Z., Friman, V. P., Gillings, M., Delgado-Baquerizo, M., & Zhu, Y. G. (2024) Prophage-encoded antibiotic resistance genes are enriched in human- impacted environments. Nat Commun 15: 8315 10.1038/s41467-024-52450-y

Liu, Y. W., Tenzintarchen, Geng, X. D., Wei, D., Dai, D. X., & Xu, R. (2020) Grazing exclusion enhanced net ecosystem carbon uptake but decreased plant nutrient content in an alpine steppe. Catena 195: 104799 10.1016/j.catena.2020.104799

Liu, Y. Z., Zhao, X. Q., Liu, W. T., Feng, B., Lv, W. D., Zhang, Z. X., Yang, X. X., & Dong, Q. M. (2024) Plant biomass partitioning in alpine meadows under different herbivores as influenced by soil bulk density and available nutrients. Catena 240: 108017 10.1016/j.catena.2024.108017

Luo, C. Y., Bao, X. Y., Wang, S. P., Zhu, X. X., Cui, S. J., Zhang, Z. H., Xu, B., Niu, H. S., Zhao, L., & Zhao, X. Q. (2015) Impacts of seasonal grazing on net ecosystem carbon exchange in alpine meadow on the Tibetan Plateau. Plant Soil 396: 381–395 10.1007/s11104-015-2602-6

Morgan, J. A., Parton, W., Derner, J. D., Gilmanov, T. G., & Smith, D. P. (2016) Importance of Early Season Conditions and Grazing on Carbon Dioxide Fluxes in Colorado Shortgrass Steppe. Rangel Ecol Manag 69: 342–350 10.1016/j.rama.2016.05.002

Nakano, T., Bavuudorj, G., Iijima, Y., & Ito, T. Y. (2020) Quantitative evaluation of grazing effect on nomadically grazed grassland ecosystems by using time-lapse cameras. Agric Ecosyst Environ 287: 106685 10.1016/j.agee.2019.106685

Niu, S. L., Wu, M. Y., Han, Y., Xia, J. Y., Li, L. H., & Wan, S. Q. (2008) Water-mediated responses of ecosystem carbon fluxes to climatic change in a temperate steppe. New Phytol 177: 209–219 10.1111/j.1469-8137.2007.02237.x

Niu, S. L., Wu, M. Y., Han, Y., Xia, J. Y., Zhang, Z., Yang, H. J., & Wan, S. Q. (2010) Nitrogen effects on net ecosystem carbon exchange in a temperate steppe. Glob Change Biol 16: 144–155 10.1111/j.1365-2486.2009.01894.x

Oesterheld, M., & McNaughton, S. J. (1991) Effect of stress and time for recovery on the amount of compensatory growth after grazing. Oecologia 85: 305–313 10.1007/BF00320604

Okach, D. O., Ondier, J. O., Kumar, A., Rambold, G., Tenhunen, J., Huwe, B., & Otieno, D. (2019) Livestock grazing and rainfall manipulation alter the patterning of CO_2_ fluxes and biomass development of the herbaceous community in a humid savanna. Plant Ecol 220: 1085–1100 10.1007/s11258-019-00977-2

Owensby, C. E., Ham, J. M., & Auen, L. M. (2006) Fluxes of CO_2_ from grazed and ungrazed tallgrass prairie. Rangel Ecol Manag 59: 111–127 10.2111/05-116r2.1

Penner, J. F., & Frank, D. A. (2021) Density-dependent plant growth drives grazer stimulation of aboveground net primary production in Yellowstone grasslands. Oecologia 196: 851–861 10.1007/s00442-021-04960-5

Pittelkow, C. M., Liang, X. Q., Linquist, B. A., van Groenigen, K. J., Lee, J., Lundy, M. E., van Gestel, N., Six, J., Venterea, R. T., & van Kessel, C. (2015) Productivity limits and potentials of the principles of conservation agriculture. Nature 517: 365–368 10.1038/nature13809

Polley, H. W., Frank, A. B., Sanabria, J., & Phillips, R. L. (2008) Interannual variability in carbon dioxide fluxes and flux-climate relationships on grazed and ungrazed northern mixed-grass prairie. Glob Change Biol 14: 1620–1632 10.1111/j.1365-2486.2008.01599.x

Qi, Y. H., Wei, D., Wang, Z. Z., Zhao, H., Fan, J. B., Tao, J., & Wang, X. D. (2024) Optimizing restoration duration to maximize CO_2_ uptake on the Tibetan Plateau. Catena 241: 108060 10.1016/j.catena.2024.108060

Reich, P. B., Wright, I. J., & Lusk, C. H. (2007) Predicting leaf physiology from simple plant and climate attributes: A global GLOPNET analysis. Ecol Appl 17: 1982–1988 10.1890/06-1803.1

Ren, S., Terrer, C., Li, J., Cao, Y. F., Yang, S. S., & Liu, D. (2024) Historical impacts of grazing on carbon stocks and climate mitigation opportunities. Nat Clim Chang 14: 380–386 10.1038/s41558-024-01957-9

Ren, S., Wang, T., Ji, X. H., Wei, L., Wei, J. J., Cao, Y. F., & Ding, J. Z. (2025) Grazing reverses climate- induced soil carbon gains on the Tibetan Plateau. Nat Commun 16: 6978 10.1038/s41467-025-62332-6

Rong, Y. P., Johnson, D. A., Wang, Z. M., & Zhua, L. L. (2017) Grazing effects on ecosystem CO_2_ fluxes regulated by interannual climate fluctuation in a temperate grassland steppe in northern China. Agric Ecosyst Environ 237: 194–202 10.1016/j.agee.2016.12.036

Shao, C. L., Chen, J. Q., & Li, L. H. (2013) Grazing alters the biophysical regulation of carbon fluxes in a desert steppe. Environ Res Lett 8: 025012 10.1088/1748-9326/8/2/025012

Shen, H., Dong, S. K., Li, S., Xiao, J. N., Han, Y. H., Yang, M. Y., Zhang, J., Gao, X. X., Xu, Y. D., Li, Y., Zhi, Y. L., Liu, S. L., Dong, Q. M., Zhou, H. K., & Yeomans, J. C. (2019) Grazing enhances plant photosynthetic capacity by altering soil nitrogen in alpine grasslands on the Qinghai-Tibetan plateau. Agric Ecosyst Environ 280: 161–168 10.1016/j.agee.2019.04.029

Shen, W., Jenerette, G. D., Hui, D., & Scott, R. L. (2016) Precipitation legacy effects on dryland ecosystem carbon fluxes: direction, magnitude and biogeochemical carryovers. Biogeosciences 13: 425–439 10.5194/bg-13-425-2016

Shi, L. A., Lin, Z. R., Tang, S. M., Peng, C. J., Yao, Z. Y., Xiao, Q., Zhou, H. K., Liu, K. S., & Shao, X. Q. (2022) Interactive effects of warming and managements on carbon fluxes in grasslands: A global meta-analysis. Agric Ecosyst Environ 340: 108178 10.1016/j.agee.2022.108178

Sloat, L. L., Gerber, J. S., Samberg, L. H., Smith, W. K., Herrero, M., Ferreira, L. G., Godde, C. M., & West, P. C. (2018) Increasing importance of precipitation variability on global livestock grazing lands. Nat Clim Chang 8: 214–218 10.1038/s41558-018-0081-5

Song, J., Wan, S., Piao, S., Knapp, A. K., Classen, A. T., Vicca, S., Ciais, P., Hovenden, M. J., Leuzinger, S., Beier, C., Kardol, P., Xia, J. Y., Liu, Q., Ru, J. Y., Zhou, Z. X., Luo, Y. Q., Guo, D. L., Langley, J. A., Zscheischler, J.,…Zheng, M. M. (2019) A meta-analysis of 1,119 manipulative experiments on terrestrial carbon-cycling responses to global change. Nat Ecol Evol 3: 1309–1320 10.1038/s41559-019-0958-3

Team, R. C. (2023) R: A language and environment for statistical computing. R Foundation for Statistical Computing

Wang, H., Li, Y. L., Zhang, J. Z., Zhang, T. R., Wang, Y. D., & Li, F. Y. (2025) Moderate grazing reduces while mowing increases greenhouse gas emissions from a steppe grassland: Key modulating function played by plant standing biomass. J Environ Manage 374: 124142 10.1016/j.jenvman.2025.124142

Wang, L., Luan, L. M., Hou, F. J., & Siddique, K. H. M. (2020) Nexus of grazing management with plant and soil properties in northern China grasslands. Sci Data 7: 39 10.1038/s41597-020-0375-0

Wang, Y. B., Zhao, Q. G., Wang, Z. W., Zhao, M. L., & Han, G. D. (2023) Overgrazing leads to decoupling of precipitation patterns and ecosystem carbon exchange in the desert steppe through changing community composition. Plant Soil 486: 607–620 10.1007/s11104-023-05894-y

Wu, Y.-S., Li, X.-R., Jia, R.-L., Yin, R.-P., & Liu, T.-J. (2023) Livestock trampling regulates the soil carbon exchange by mediating surface roughness and biocrust cover. Geoderma 429: 116275 10.1016/j.geoderma.2022.116275

Wu, Y. T., Li, H., Cui, J. H., Han, Y., Li, H. Y., Miao, B. L., Tang, Y. K., Li, Z. Y., Zhang, J. H., Wang, L. X., & Liang, C. Z. (2024) Precipitation variation: a key factor regulating plant diversity in semi-arid livestock grazing lands. Front Plant Sci 15: 1294895 10.3389/fpls.2024.1294895

Xiao, J. F., Zhuang, Q. L., Law, B. E., Baldocchi, D. D., Chen, J. Q., Richardson, A. D., Melillo, J. M., Davis, K. J., Hollinger, D. Y., Wharton, S., Oren, R., Noormets, A., Fischer, M. L., Verma, S. B., Cook, D. R., Sun, G., McNulty, S., Wofsy, S. C., Bolstad, P. V.,…Torn, M. S. (2011) Assessing net ecosystem carbon exchange of US terrestrial ecosystems by integrating eddy covariance flux measurements and satellite observations. Agric For Meteorol 151: 60–69 10.1016/j.agrformet.2010.09.002

Xun, W., Yan, R., Ren, Y., Jin, D., Xiong, W., Zhang, G., Cui, Z., Xin, X., & Zhang, R. (2018) Grazing- induced microbiome alterations drive soil organic carbon turnover and productivity in meadow steppe. Microbiome 6: 170 10.1186/s40168-018-0544-y

Yang, N., Zohner, C. M., Crowther, T. W., Feng, J. G., Wu, J., Chen, X. L., Han, W. X., Stocker, B. D., Hui, D. F., Augusto, L., Yue, K., Hou, E. Q., Jiang, M. K., Feng, H. L., Chen, Z. X., Wu, W. J., Xing, A. J., Chen, C. R., Sardans, J.,…Yan, Z. B. (2025) Leaf economic strategies drive global variation in phosphorus stimulation of terrestrial plant production. Nat Commun 16: 5562 10.1038/s41467-025-60633-4

Yin, M. Y., Gao, X. P., Kuang, W. N., & Tenuta, M. (2023) Soil N_2_O emissions and functional genes in response to grazing grassland with livestock: A meta-analysis. Geoderma 436 10.1016/j.geoderma.2023.116538

Yu, H. L., Wang, X., Wu, Y. Q., Wang, C. W., Yan, R. R., Xu, D. W., Yan, Y. C., & Xin, X. P. (2025) Light grazing tends to enhance ecosystem carbon sequestration and resource use efficiency in a meadow steppe of northern China. Agric For Meteorol 372: 110690 10.1016/j.agrformet.2025.110690

Zhang, R. Y., Tian, D. S., Chen, H. Y. H., Seabloom, E. W., Han, G. D., Wang, S. P., Yu, G. R., Li, Z. L., & Niu, S. L. (2022) Biodiversity alleviates the decrease of grassland multifunctionality under grazing disturbance: A global meta-analysis. Glob Ecol Biogeogr 31: 155–167 10.1111/geb.13408

Zhang, S., Chen, W., Wang, Y. N., Li, Q., Shi, H. M., Li, M., Sun, Z. X., Zhu, B. R., & Seyoum, G. (2024) Human interventions have enhanced the net ecosystem productivity of farmland in China. Nat Commun 15: 10523 10.1038/s41467-024-54907-6

Zhou, G. Y., Luo, Q., Chen, Y. J., Hu, J. Q., He, M., Gao, J., Zhou, L. Y., Liu, H. Y., & Zhou, X. H. (2019) Interactive effects of grazing and global change factors on soil and ecosystem respiration in grassland ecosystems: A global synthesis. J Appl Ecol 56: 2007–2019 10.1111/1365-2664.13443

Zhou, G. Y., Zhou, X. H., He, Y. H., Shao, J. J., Hu, Z. H., Liu, R. Q., Zhou, H. M., & Hosseinibai, S. (2017) Grazing intensity significantly affects belowground carbon and nitrogen cycling in grassland ecosystems: a meta-analysis. Glob Change Biol 23: 1167–1179 10.1111/gcb.13431

Zhou, X., Wan, S., & Luo, Y. (2007) Source components and interannual variability of soil CO_2_ efflux under experimental warming and clipping in a grassland ecosystem. Glob Change Biol 13: 761–775 10.1111/j.1365-2486.2007.01333.x

Zhou, Y. P., Xu, M., Ren, S., Du, Y. X., Yue, Y. H., Yu, H. R., Zhang, Y., Jiang, S. C., Xu, T. T., & Wang, L. (2025) More Than a Decade of Moderate Grazing: No Impact on Soil Organic Carbon Stocks and Enhancement of Mineral-Associated Organic Carbon via Livestock Diversification. Glob Change Biol 31: e70466 10.1111/gcb.70466

Zhu, L. L., Johnson, D. A., Wang, W. G., Ma, L., & Rong, Y. P. (2015) Grazing effects on carbon fluxes in a Northern China grassland. J Arid Environ 114: 41–48 10.1016/j.jaridenv.2014.11.004

Zong, N., & Shi, P. L. (2019) Nitrogen addition stimulated compensatory growth responses to clipping defoliation in a Northern Tibetan alpine meadow. Grassl Sci 65: 60–68 10.1111/grs.12219

