## Supplementary materials for "Wetness modulates the effects of grazing on net ecosystem productivity in global grasslands"

**Table** S1 Summary of meta-analyses on the effects of grazing intensity on ecosystem CO_2_ fluxes

| Region | GPP | | | | ER | | | | NEP | | | |
| --- | --- | --- | --- | --- | --- | --- | --- | --- | --- | --- | --- | --- |
|  | Grazing | LG | MG | HG | Grazing | LG | MG | HG | Grazing | LG | MG | HG |
| The Tibetan Plateau, China (Shi et al., 2022) | ns | - | - | - | ns | - | - | - | ns | - | - | - |
| Global grasslands (Wan et al., 2025) | - | - | - | - | ↑ | - | - | - | ns | - | - | - |
| Global grasslands (Zhou et al., 2019) | - | - | - | - | ↑ | - | - | - | - | - | - | - |
| Global grasslands (Zhang et al., 2022) | - | ↓ | ↓ | ↓ | - | ↓ | ↓ | ↓ | - | ↓ | ↓ | ↓ |
| Chinese grasslands (Jiang et al., 2020) | - | - | - | - | ↓ | ↓ | ↓ | ↓ | ↑ | ↑ | ↑ | ↑ |
| Global grasslands in this study | ↓ | ↓ | ↓ | ↓ | ↓ | ↓ | ↓ | ↓ | ↓ | ↓ | ns | ↓ |

Note: GPP: gross primary productivity; ER: ecosystem respiration; NEP: net ecosystem productivity; LG, light grazing; MG, moderate grazing; HG, heavy grazing; ↓: decrease; ↑ increase; ns: not significant.

**Table S2** Effect of grazing intensities on mean ecosystem CO_2_ fluxes and plant biomass

| Variable | Treatment | Mean | SD | SE |  | *df* | Sum Sq | Mean Sq | *F* value | *P* value |
| --- | --- | --- | --- | --- | --- | --- | --- | --- | --- | --- |
| Gross primary productivity | CK | 9.91 | 0.71 | 0.41 | **Treatment** | **3** | **20.519** | **6.84** | **18.48** | **<0.001^***^** |
|  | LG | 10.14 | 0.56 | 0.33 |  |  |  |  |  |  |
|  | MG | 8.18 | 0.24 | 0.14 | Residuals | 8 | 2.961 | 0.37 |  |  |
|  | HG | 6.95 | 0.77 | 0.45 |  |  |  |  |  |  |
| Ecosystem respiration | CK | 7.66 | 0.33 | 0.19 | **Treatment** | **3** | **12.565** | **4.188** | **65.8** | **<0.001^***^** |
|  | LG | 7.54 | 0.12 | 0.07 |  |  |  |  |  |  |
|  | MG | 5.92 | 0.15 | 0.09 | Residuals | 8 | 0.509 | 0.064 |  |  |
|  | HG | 5.29 | 0.32 | 0.19 |  |  |  |  |  |  |
| Net ecosystem productivity | CK | 2.36 | 0.47 | 0.27 | **Treatment** | **3** | **1.851** | **0.617** | **3.611** | **0.07** |
|  | LG | 2.94 | 0.28 | 0.16 |  |  |  |  |  |  |
|  | MG | 2.27 | 0.35 | 0.20 | Residuals | 8 | 1.367 | 0.1709 |  |  |
|  | HG | 1.84 | 0.51 | 0.29 |  |  |  |  |  |  |
| Aboveground biomass | CK | 181.07 | 13.11 | 7.57 | **Treatment** | **3** | **7969** | **2656.4** | **11.16** | **0.01^**^** |
|  | LG | 157.48 | 15.12 | 8.73 |  |  |  |  |  |  |
|  | MG | 156.12 | 22.62 | 13.06 | Residuals | 8 | 1903 | 237.9 |  |  |
|  | HG | 109.97 | 6.26 | 3.61 |  |  |  |  |  |  |
| Belowground biomass | CK | 1219.77 | 112.31 | 64.84 | **Treatment** | **3** | **88887** | **29629** | **3.033** | **0.09** |
|  | LG | 1119.46 | 145.11 | 83.78 |  |  |  |  |  |  |
|  | MG | 1065.90 | 67.81 | 39.15 | Residuals | 8 | 78162 | 9770 |  |  |
|  | HG | 982.61 | 28.49 | 16.45 |  |  |  |  |  |  |
| Root/Shoot ratio | CK | 8.53 | 1.06 | 0.61 | **Treatment** | **3** | **13.089** | **4.363** | **4.133** | **<0.05^*^** |
|  | LG | 8.71 | 1.35 | 0.78 |  |  |  |  |  |  |
|  | MG | 9.09 | 0.98 | 0.57 | Residuals | 8 | 8.445 | 1.056 |  |  |
|  | HG | 11.14 | 0.56 | 0.33 |  |  |  |  |  |  |

Note: CK, without grazing; LG, light grazing; MG, moderate grazing, HG, heavy grazing. ^*^ *P* < 0.05; ^**^ *P* < 0.001; ^***^ *P* < 0.0001. **Table S3** The ordinary least squares linear regression models of ecosystem CO_2_ fluxes response to wetness index, soil temperature, soil moisture, and of plant biomass in response to soil temperature and grazing duration under different grazing intensities

| Predictors | Grazing  intensity | Model Parameters (Estimate ± SE) | | *n* | *df1* | *df2* | *R_adj_*^2^ | *P* |
| --- | --- | --- | --- | --- | --- | --- | --- | --- |
|  |  | Intercept | Slope |  |  |  |  |  |
| Gross primary productivity & Wetness index | **Overall** | **-0.775 ± 0.174 ^***^** | **0.026 ± 0.008 ^**^** | **45** | **1** | **43** | **0.180** | **0.002** |
|  | **LG** | **-0.939 ± 0.132 ^***^** | **0.043 ± 0.006 ^***^** | **15** | **1** | **13** | **0.780** | **<0.001** |
|  | MG | -0.428 ± 0.156 **^*^** | 0.010 ± 0.007 | 15 | 1 | 13 | 0.072 | 0.172 |
|  | HG | -0.958 ± 0.341 **^*^** | 0.025 ± 0.016 | 15 | 1 | 13 | 0.095 | 0.139 |
| Net ecosystem productivity & Wetness index | **Overall** | **-2.049 ± 0.452 ^***^** | **0.082 ±0.021** | **45** | **1** | **43** | **0.249** | **<0.001** |
|  | **LG** | **-1.992 ± 0.442 ^***^** | **0.095 ± 0.020 ^***^** | **15** | **1** | **13** | **0.599** | **<0.001** |
|  | MG | -1.606 ± 0.702 ^*^ | 0.063 ± 0.032 | 15 | 1 | 13 | 0.171 | 0.070 |
|  | HG | -2.552 ± 0.969 ^*^ | 0.088 ± 0.045 | 15 | 1 | 13 | 0.169 | 0.072 |
| Gross primary productivity & Soil moisture | **Overall** | **-0.189 ± 0.044 ^***^** | **0.531 ± 0.219 ^*^** | **45** | **1** | **43** | **0.100** | **0.019** |
|  | LG | -0.026 ± 0.069 | -0.016 ± 0.511 | 15 | 1 | 13 | -0.077 | 0.975 |
|  | MG | -0.207 ± 0.038 ^***^ | -0.095 ± 0.192 | 15 | 1 | 13 | -0.057 | 0.631 |
|  | **HG** | **-0.257 ± 0.108 ^*^** | **0.952 ± 0.426 ^*^** | **15** | **1** | **13** | **0.222** | **0.043** |
| Gross primary productivity &  Soil temperature | Overall | -0.203 ± 0.045 ^***^ | -0.491 ± 0.297 | 45 | 1 | 43 | 0.038 | 0.106 |
|  | LG | -0.024 ± 0.066 | -0.431 ± 0.476 | 15 | 1 | 13 | -0.013 | 0.381 |
|  | MG | -0.209 ± 0.039 ^***^ | 0.003 ± 0.261 | 15 | 1 | 13 | -0.077 | 0.989 |
|  | HG | -0.435 ± 0.118 ^**^ | -0.006 ± 0.704 | 15 | 1 | 13 | -0.077 | 0.993 |
| Net ecosystem productivity & Soil temperature | Overall | -0.256 ± -0.123 | -1.407 ± 0.806 | 45 | 1 | 43 | 0.044 | 0.088 |
|  | LG | 0.022 ± 0.168 | -0.293 ± 1.213 | 15 | 1 | 13 | -0.072 | 0.813 |
|  | MG | -0.253 ± 0.175 | -1.459 ± 1.171 | 15 | 1 | 13 | 0.038 | 0.235 |
|  | HG | -0.627 ± 0.348 | -0.649 ± 2.083 | 15 | 1 | 13 | -0.069 | 0.759 |
| Aboveground biomass & Soil temperature | **Overall** | **-0.274 ± 0.040 ^***^** | **-0.834 ± 0.265** | **45** | **1** | **43** | **0.168** | **0.003** |
|  | **LG** | **-0.172 ± 0.039 ^***^** | **-1.041 ± 0.287 ^**^** | **15** | **1** | **13** | **0.465** | **0.003** |
|  | MG | -0.186 ± 0.059 ^**^ | -0.512 ± 0.392 | 15 | 1 | 13 | 0.048 | 0.214 |
|  | HG | -0.597 ± 0.087 ^***^ | 0.338 ± 0.518 | 15 | 1 | 13 | -0.043 | 0.525 |
| Aboveground biomass & Duration | Overall | -0.554 ± 0.275 | 0.027 ± 0.030 | 45 | 1 | 43 | -0.004 | 0.370 |
|  | LG | -0.054 ± 0.360 | -0.013 ±0.039 | 15 | 1 | 13 | -0.067 | 0.738 |
|  | MG | -0.006 ± 0.399 | -0.020 ±0.044 | 15 | 1 | 13 | -0.059 | 0.649 |
|  | **HG** | **-1.603 ± 0.294 ^***^** | **0.116 ±0.032 ^**^** | **15** | **1** | **13** | **0.459** | **0.003** |
| Belowground biomass & Duration | Overall | 0.014 ± 0.163 | -0.017 ± 0.018 | 45 | 1 | 43 | -0.001 | 0.337 |
|  | LG | 0.045 ± 0.356 | -0.016 ± 0.039 | 15 | 1 | 13 | -0.064 | 0.696 |
|  | MG | 0.040 ± 0.234 | -0.019 ± 0.026 | 15 | 1 | 13 | -0.035 | 0.482 |
|  | HG | -0.042 ± 0.252 | -0.018 ± 0.028 | 15 | 1 | 13 | -0.042 | 0.522 |

Note: Overall, overall grazing included light, moderate and heavy grazing; CK, without grazing; LG, light grazing; MG, moderate grazing, HG, heavy grazing. ^**^ *P* < 0.001; ^***^ *P* < 0.0001.

**Table S4** Results from Egger's tests of publication bias on our models

|  | *z* | *P* |
| --- | --- | --- |
| Gross primary productivity | 0.57 | 0.567 |
| Ecosystem respiration | 0.98 | 0.322 |
| Net ecosystem productivity | 0.97 | 0.329 |
| Aboveground biomass | -0.72 | 0.469 |
| Belowground biomass | -0.38 | 0.697 |
| Root/Shoot ratio | -1.01 | 0.311 |
| Soil moisture | 0.08 | 0.93 |
| Soil temperature | 0.09 | 0.92 |

Note: The *P*-value above 0.05 suggests there is no publication bias.

**Fig.** S1 The monthly total precipitation and average air temperature (A), annual precipitation (B) and wetness index (C) in the study area of the grazing intensity experiment in the typical steppe from 2019 to 2023.

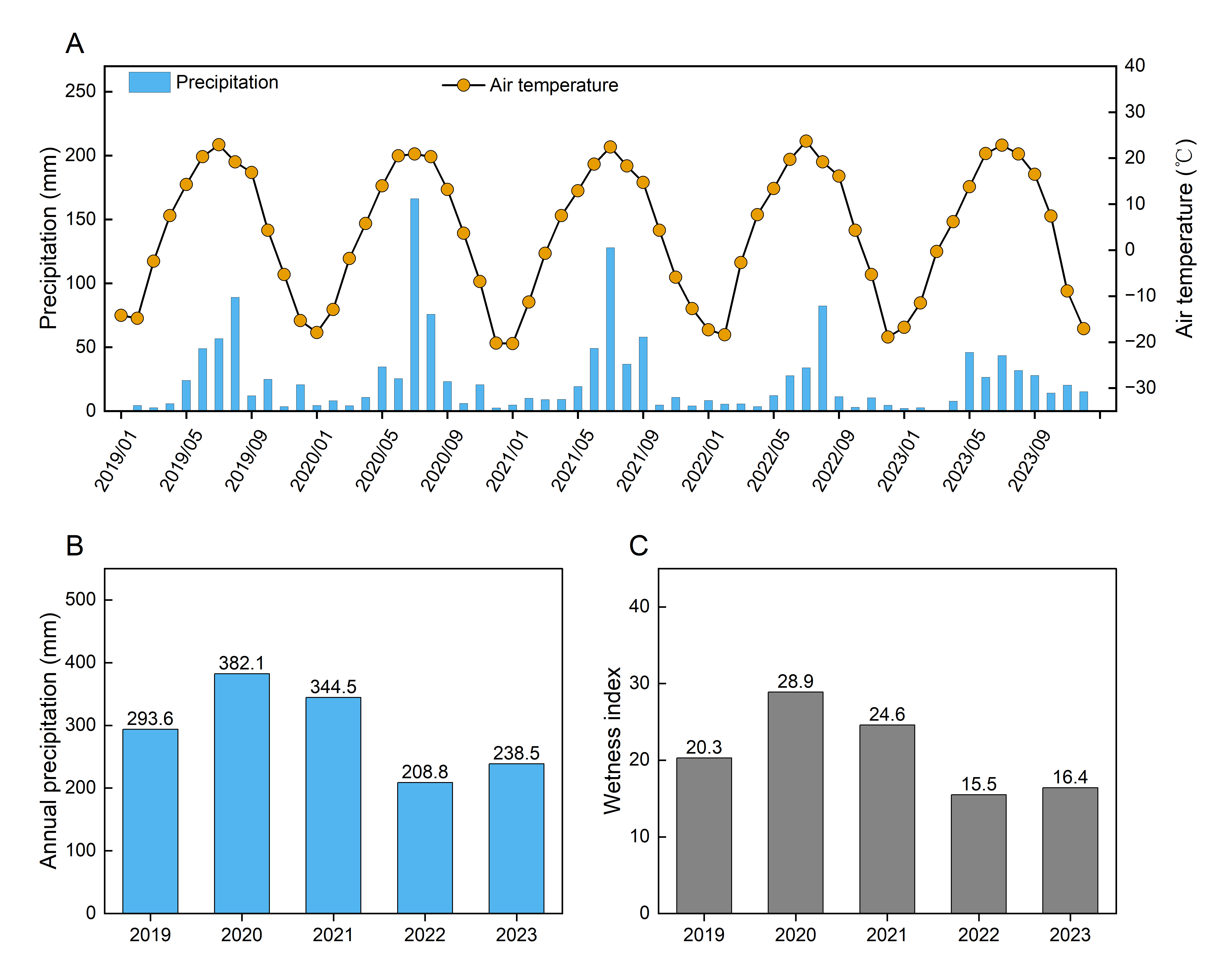

**Fig.** S2 The design and experimental plots of the grazing intensity experiment in the typical steppe, Inner Mongolia, China (A-B). The figures of net ecosystem productivity (NEP) and ecosystem respiration (ER) being measured (C). CK, without grazing; LG, light grazing; MG, moderate grazing; HG, heavy grazing.

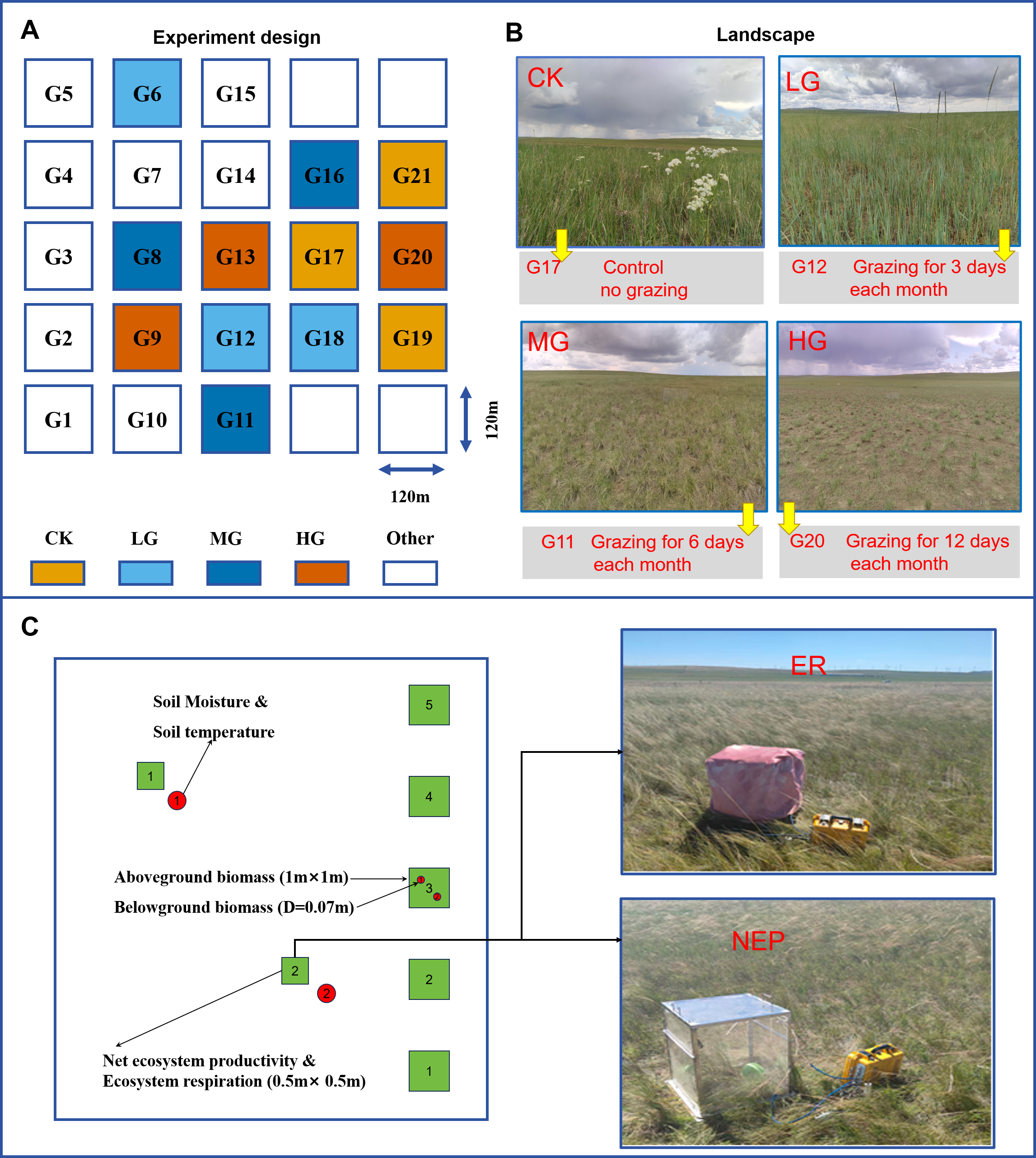

**Fig.** S3 The random forest model of relative importance of environmental predictors of gross primary productivity (A), ecosystem respiration (B), net ecosystem productivity (C), aboveground biomass (D) and belowground biomass (E) in the field experiment from 2019 to 2023. ^*^*P* < 0.05, ^**^*P* < 0.01 and ^***^*P* < 0.001. Soil moisture_RR, the response ratio of soil moisture; Soil temperature_RR, the response ratio of soil temperature.

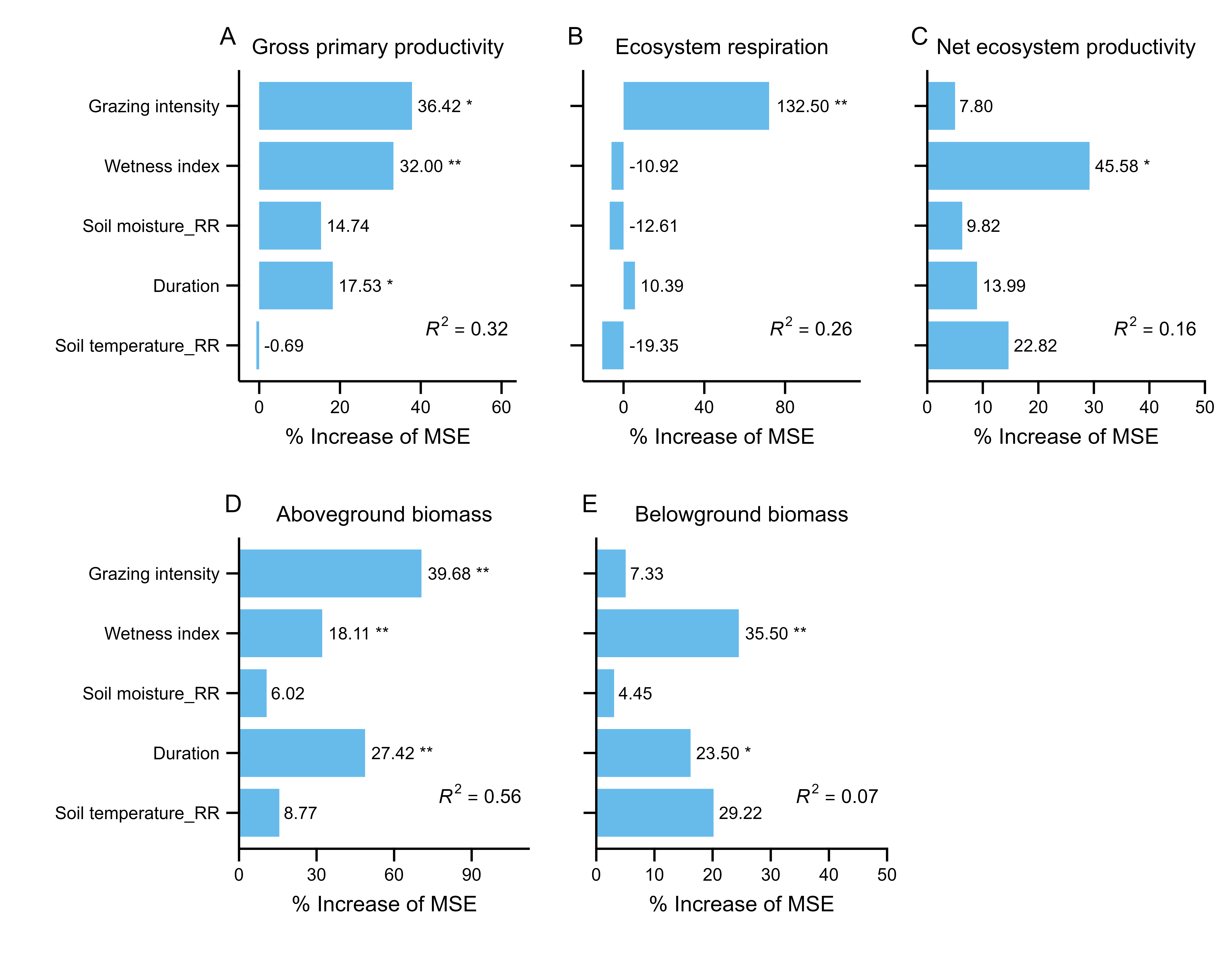

**Fig.** S4 The effects of grazing on aboveground net primary productivity (ANPP) (A) and relative growth rate (B) and their relationships with net ecosystem productivity (NEP) (C-D) in the field experiment from 2019 to 2023. Data (means ± SE, *n* = 3) followed by different lowercase letters indicate significant differences between treatments at *P* < 0.05. The significant regression lines and 95% confidence intervals are shown with lines (solid for significant) and shaded areas, respectively. CK, without grazing; LG, light grazing; MG, moderate grazing; HG, heavy grazing.

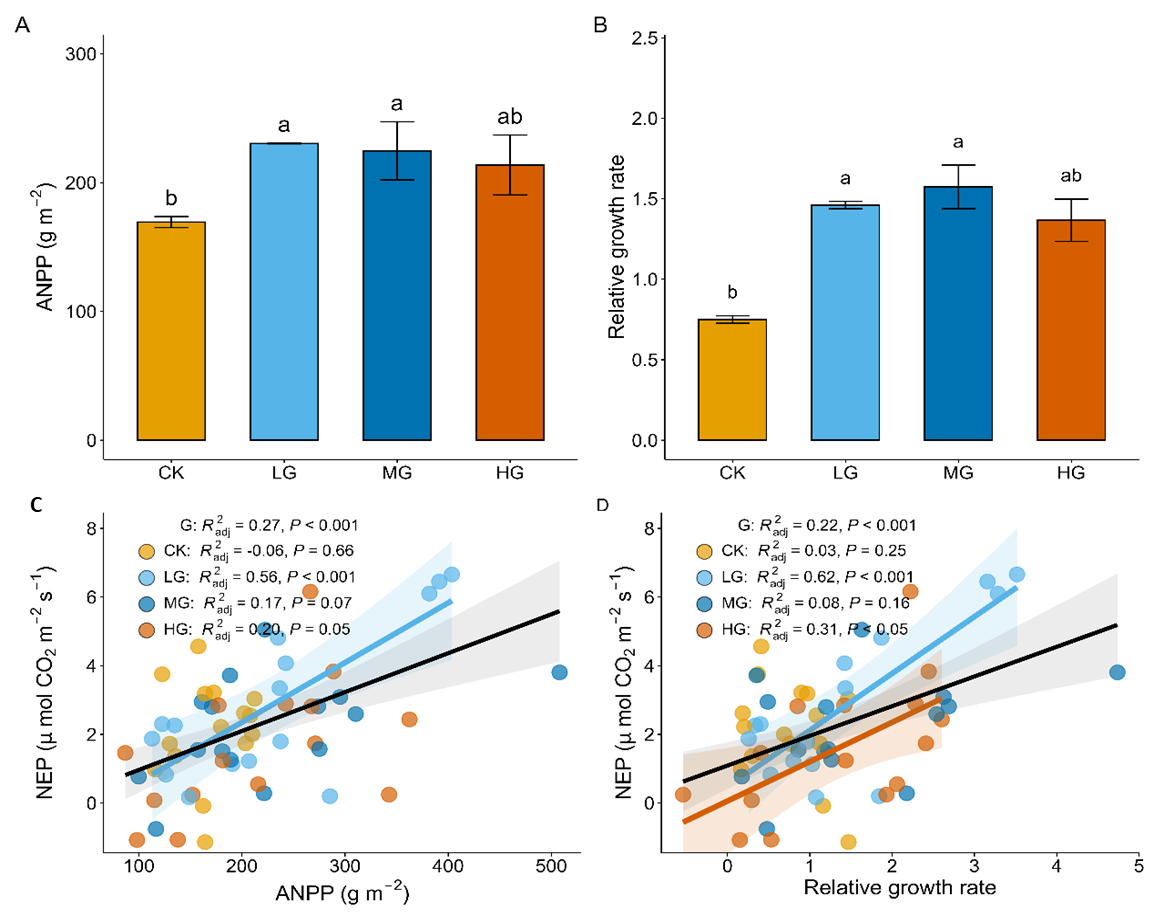

**Fig.** S5 The mean relative growth rate in different grazing intensity treatments from 2019 to 2023 (A), relationships between the relative growth rate and mean annual precipitation and wetness index, respectively (B-C). The significant regression lines and 95% confidence intervals are shown with lines and shaded areas, respectively. CK, without grazing; LG, light grazing; MG, moderate grazing; HG, heavy grazing. Data (means ± SE, *n* = 3) followed by different lowercase and uppercase letters indicate significant differences between treatments and years, respectively, at *P* < 0.05.

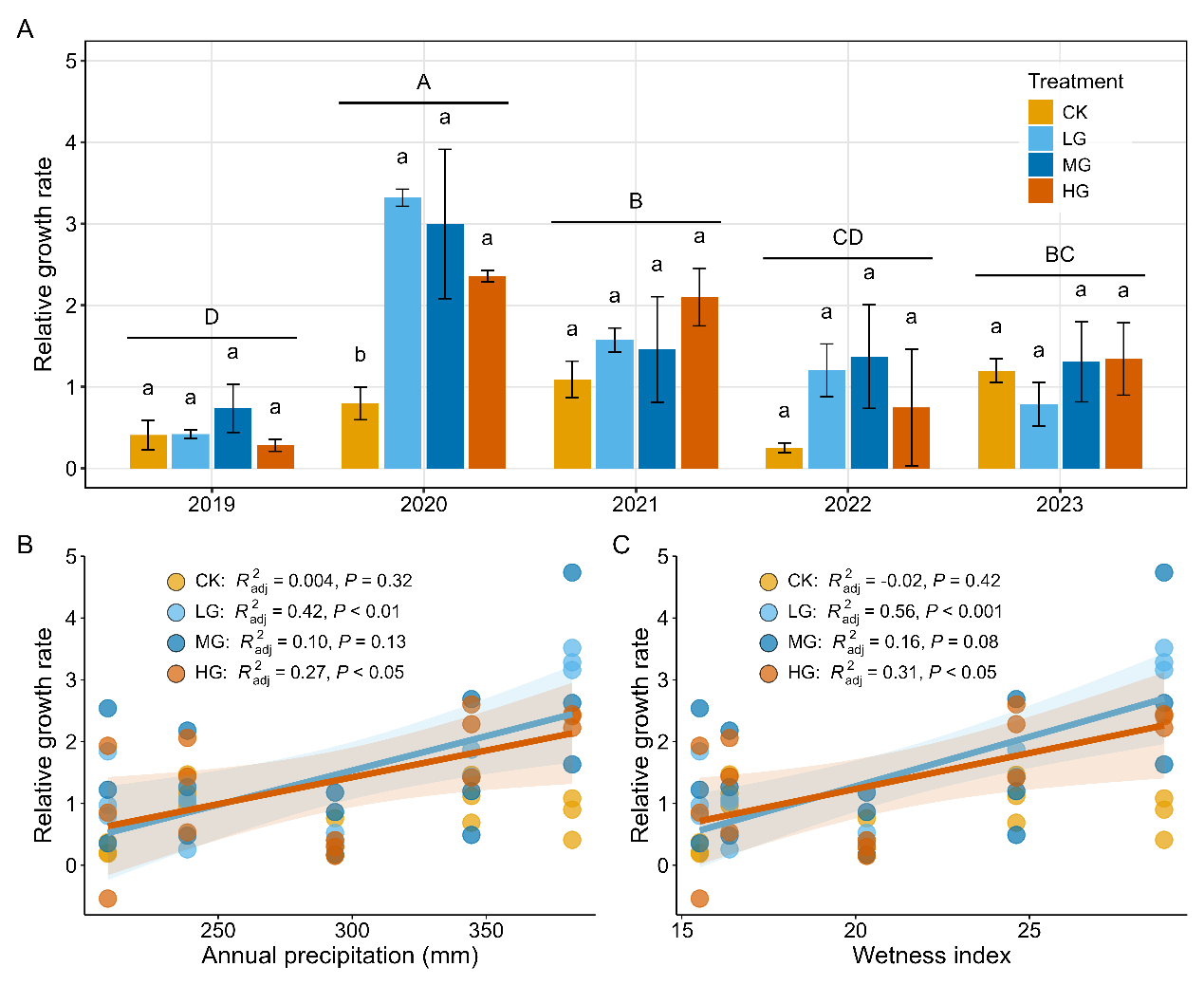

**Fig.** S6 Relationships between ecosystem CO_2_ fluxes and wetness index. The significant regression lines and 95% confidence intervals are shown with lines and shaded areas, respectively. CK, without grazing; LG, light grazing; MG, moderate grazing; HG, heavy grazing.

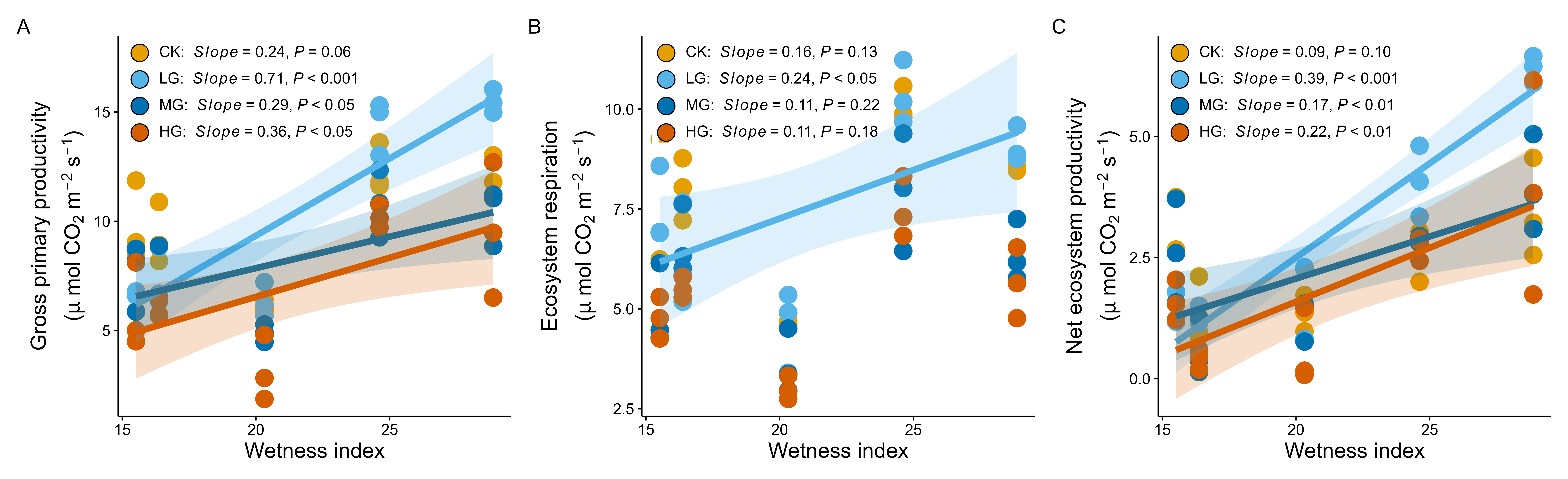

**Fig.** S7 Effects of different grazing intensities on C_3_, C_4_ plant biomass, C_3_:C_4_ plant biomass ratio (A-C), C_3_, C_4_ plant richness, and C_3_:C_4_ plant richness ratio (D-F) in the field experiment of the typical steppe. CK, without grazing; LG, light grazing; MG, moderate grazing; HG, heavy grazing. Data (means ± SE, *n* = 3) followed by different lowercase letters indicate differences at *P* < 0.05 between treatments.

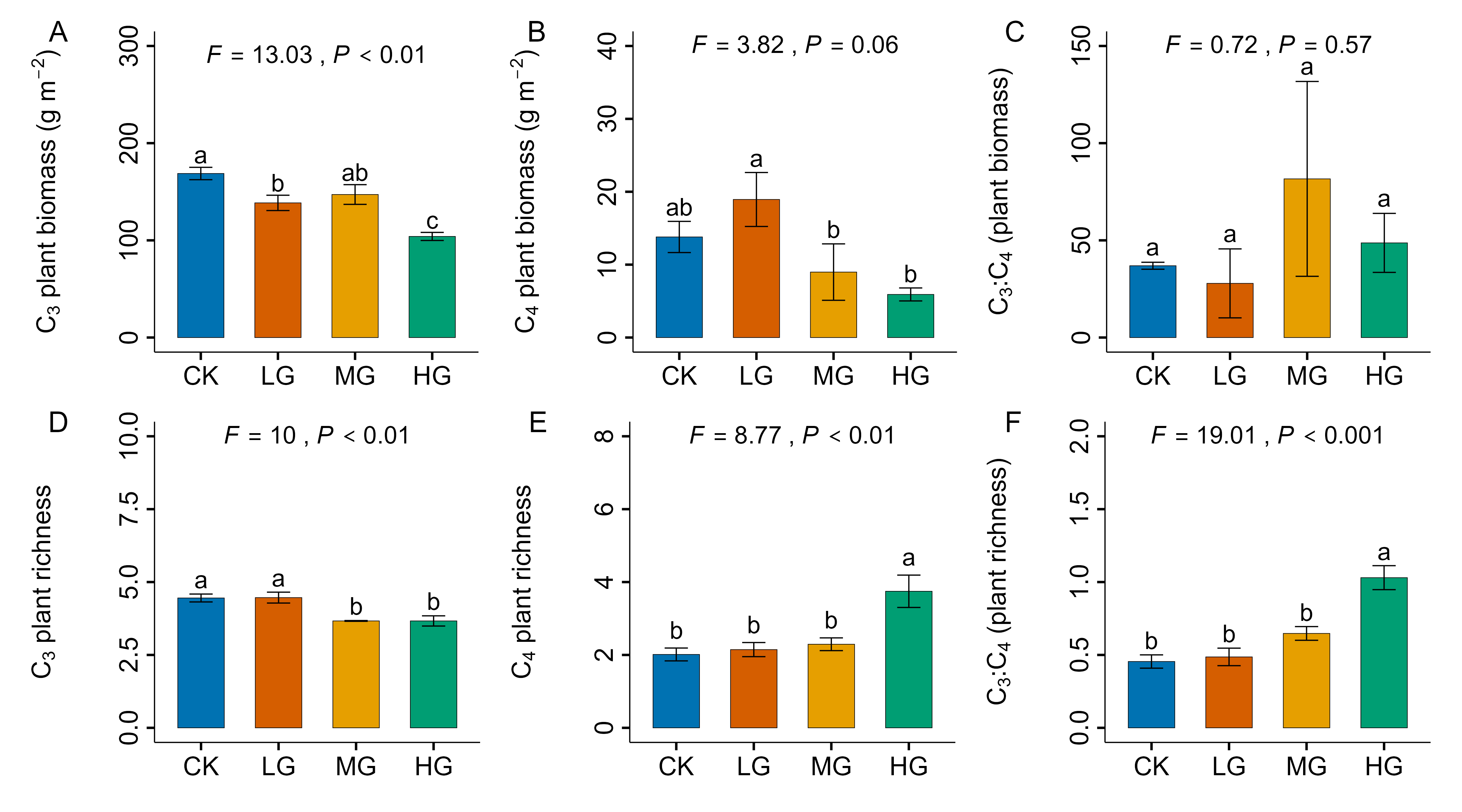

**Fig.** S8 Hypothesized structural equation models (SEMs) (A-B) and the fitted models (C-D) for the effect of grazing intensity and wetness index on net ecosystem productivity (NEP). We hypothesized that grazing intensity and wetness index may influence NEP directly or indirectly through changes in soil temperature (ST), soil moisture (SM), relative growth rate (RGR), aboveground biomass (AGB), and C_3_ and C_4_ plant functional group composition. Panels A-C represented alternative candidate models, and panel D showed the final supported SEM. Arrows indicated hypothesized relationships among variables. Significant pathways in the final model were shown with standardized path coefficients.

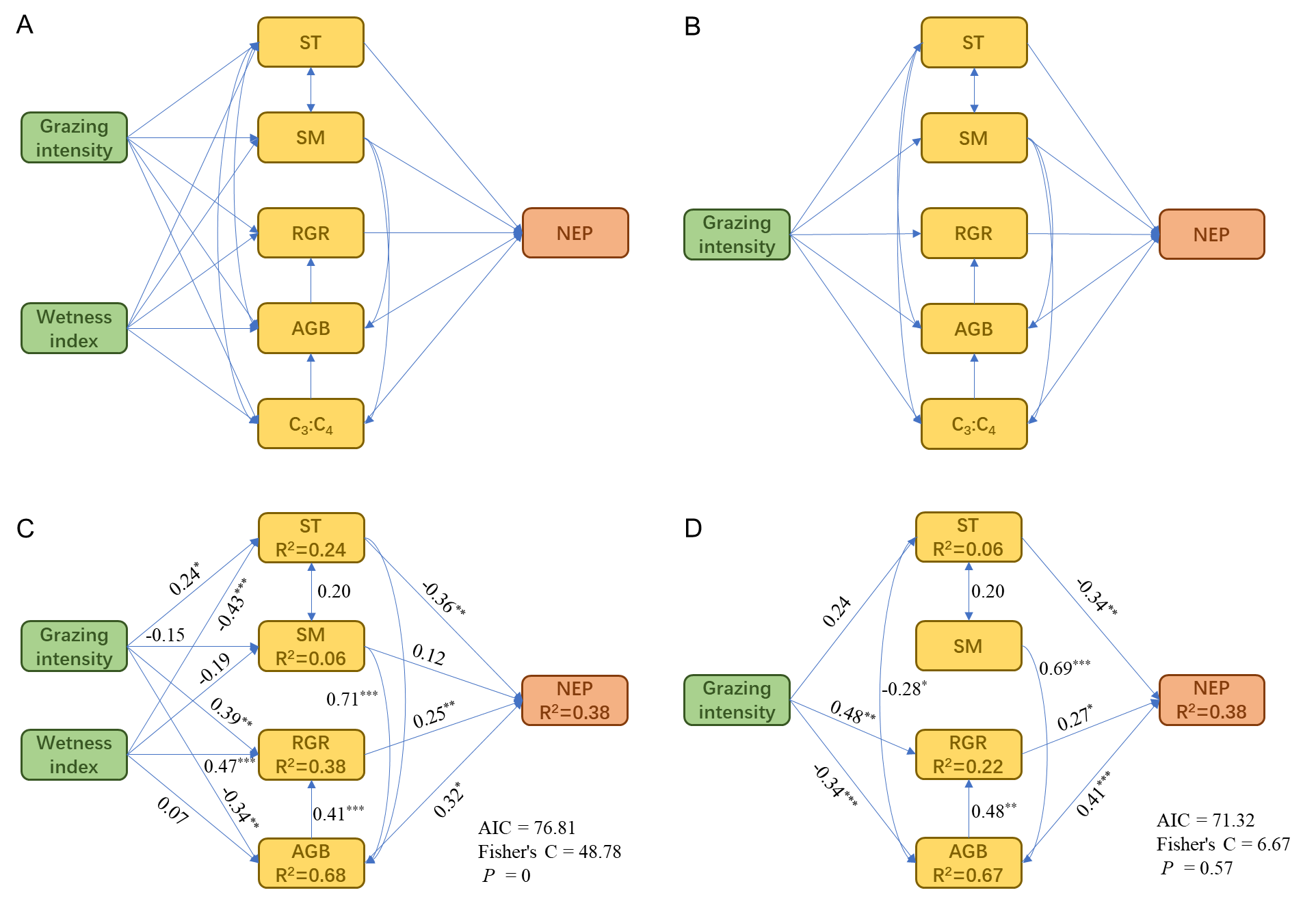
**Fig.** S9 PRISMA flow diagram of the meta-analysis in this study.

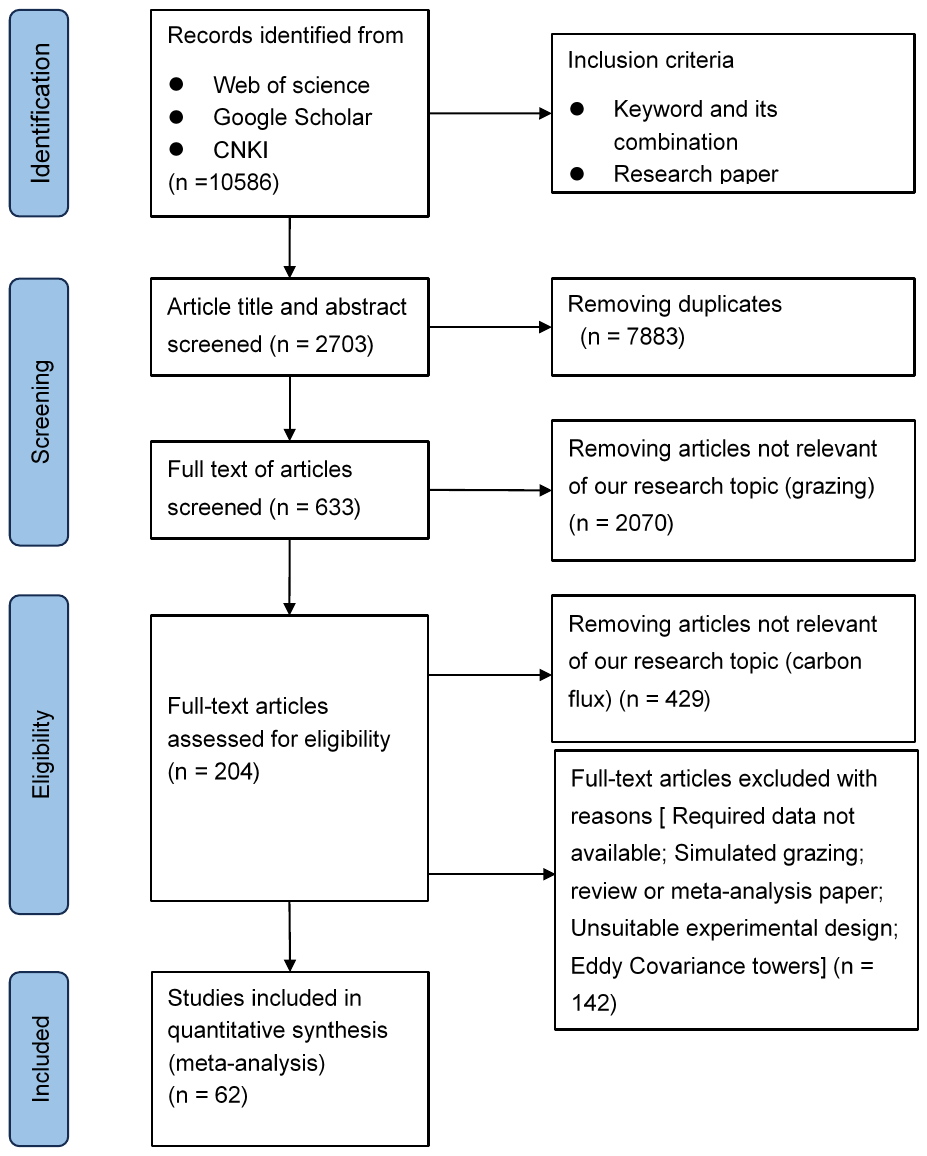

**Fig.** S10 Responses of the individual effects of grazing on ecosystem CO_2_ fluxes at different levels of mean annual precipitation (A), mean annual air temperature (B) and grazing duration (C) in global grasslands from the meta-analysis. Circles and error bars represent average parameter estimates and 95% confidence interval (CI). The sample size of observations for each variable is shown on the left.

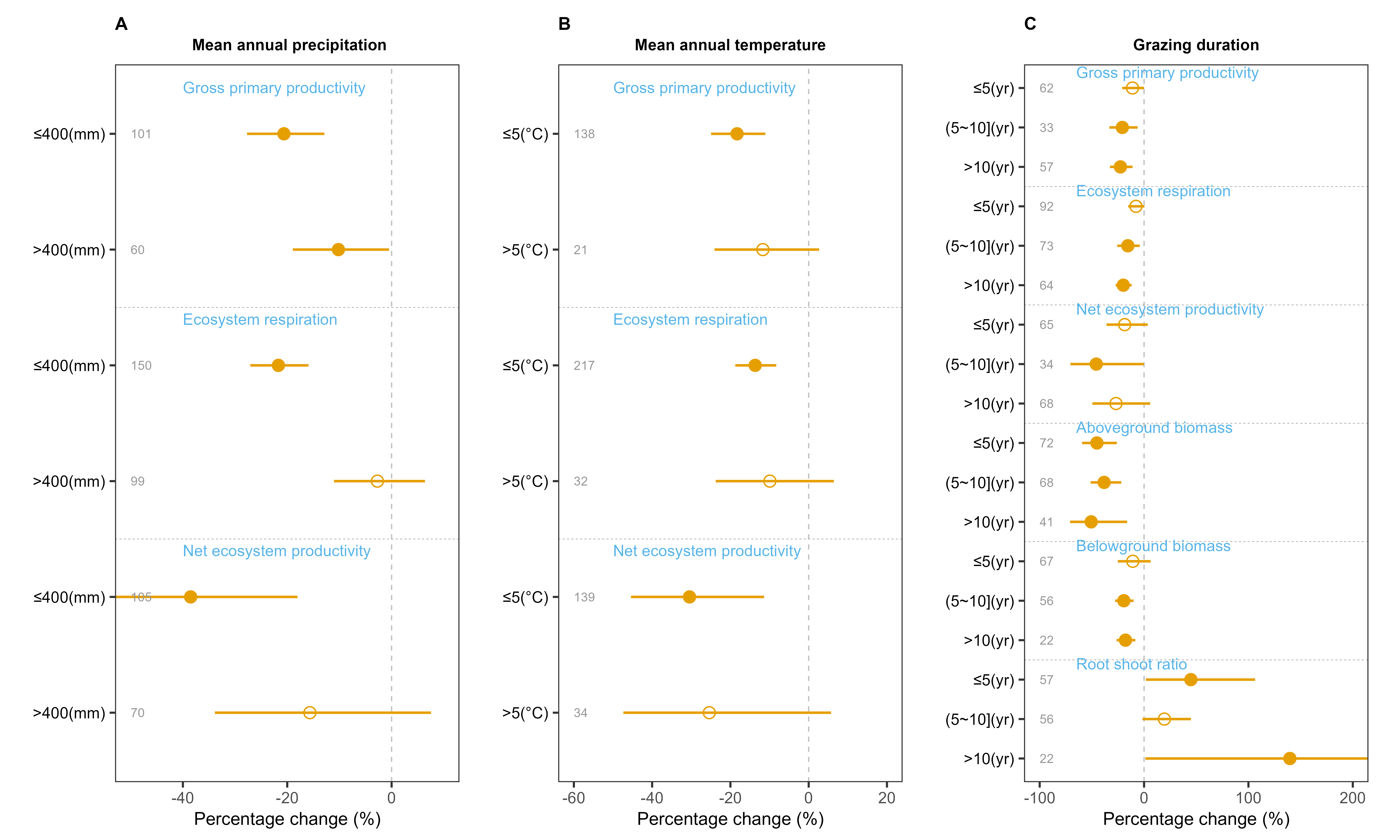

**Fig.** S11 Responses of the individual effects of grazing on ecosystem CO_2_ fluxes (gross primary productivity (A), ecosystem respiration (B), and net ecosystem productivity (C) at different levels of grassland type (desert grassland, temperate grassland, alpine grassland, savanna, and others) in global grasslands in the meta-analysis. Circles and error bars represent average parameter estimates and 95% confidence interval (CI). The sample size of observations for each variable is shown on the left.

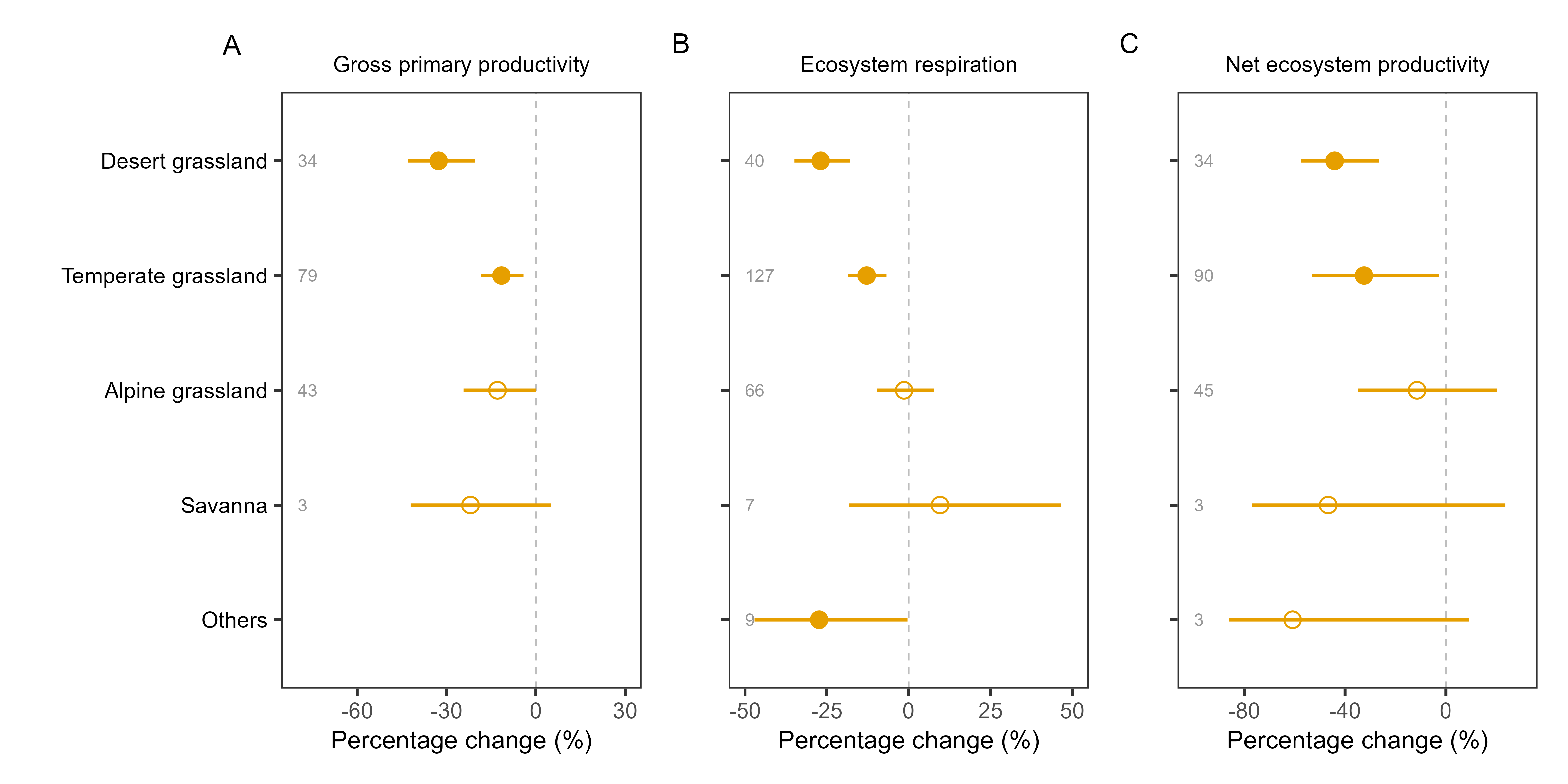

**Fig.** S12 Responses of the individual effects of grazing on ecosystem CO_2_ fluxes (gross primary productivity (A), ecosystem respiration (B), and net ecosystem productivity (C)) at different livestock types (cattle, sheep, and mix) in global grasslands of the meta-analysis. Circles and error bars represent average parameter estimates and 95% confidence interval (CI). The sample size of observations for each variable is shown on the left.

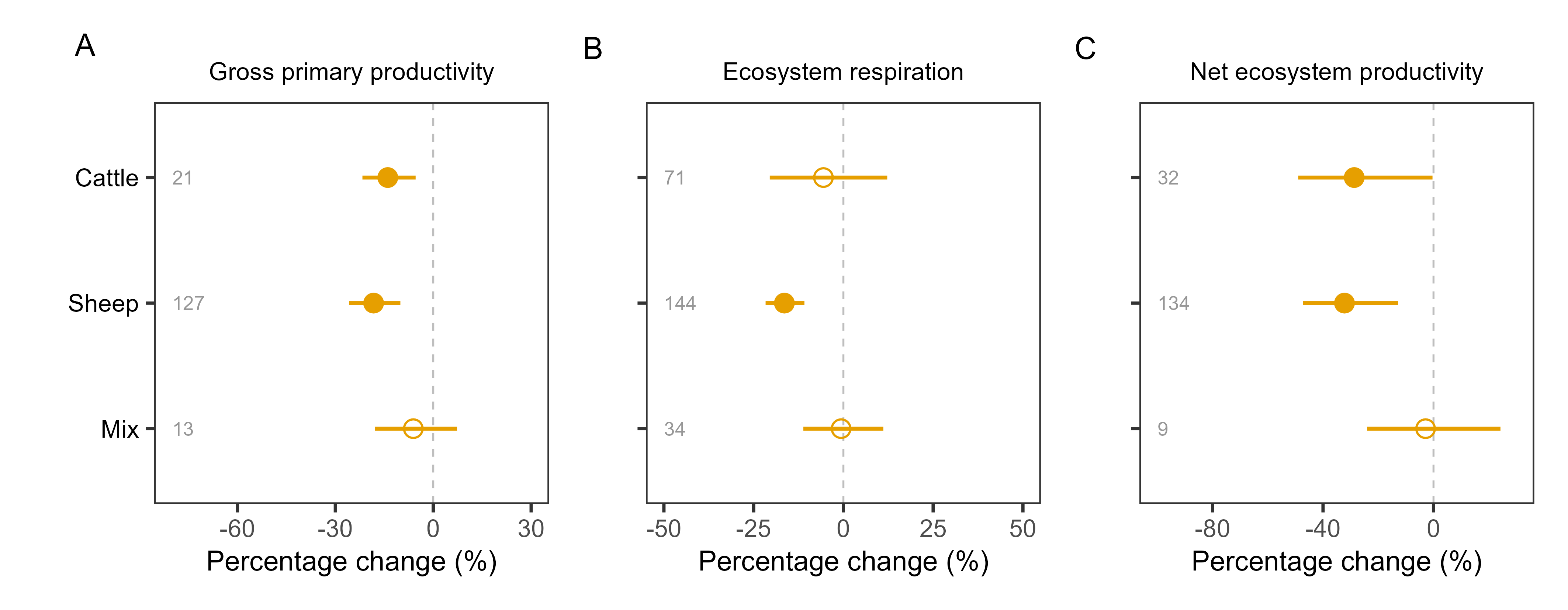

**Fig.** S13 The random forest model of the relative importance of environmental predictors of gross primary productivity (A), ecosystem respiration (B), net ecosystem productivity (C), aboveground biomass (D) and belowground biomass (E) in global grasslands of the meta-analysis. ^*^*P* < 0.05, ^**^*P* < 0.01. Soil moisture_RR, the response ratio of soil moisture; Soil temperature_RR, the response ratio of soil temperature.

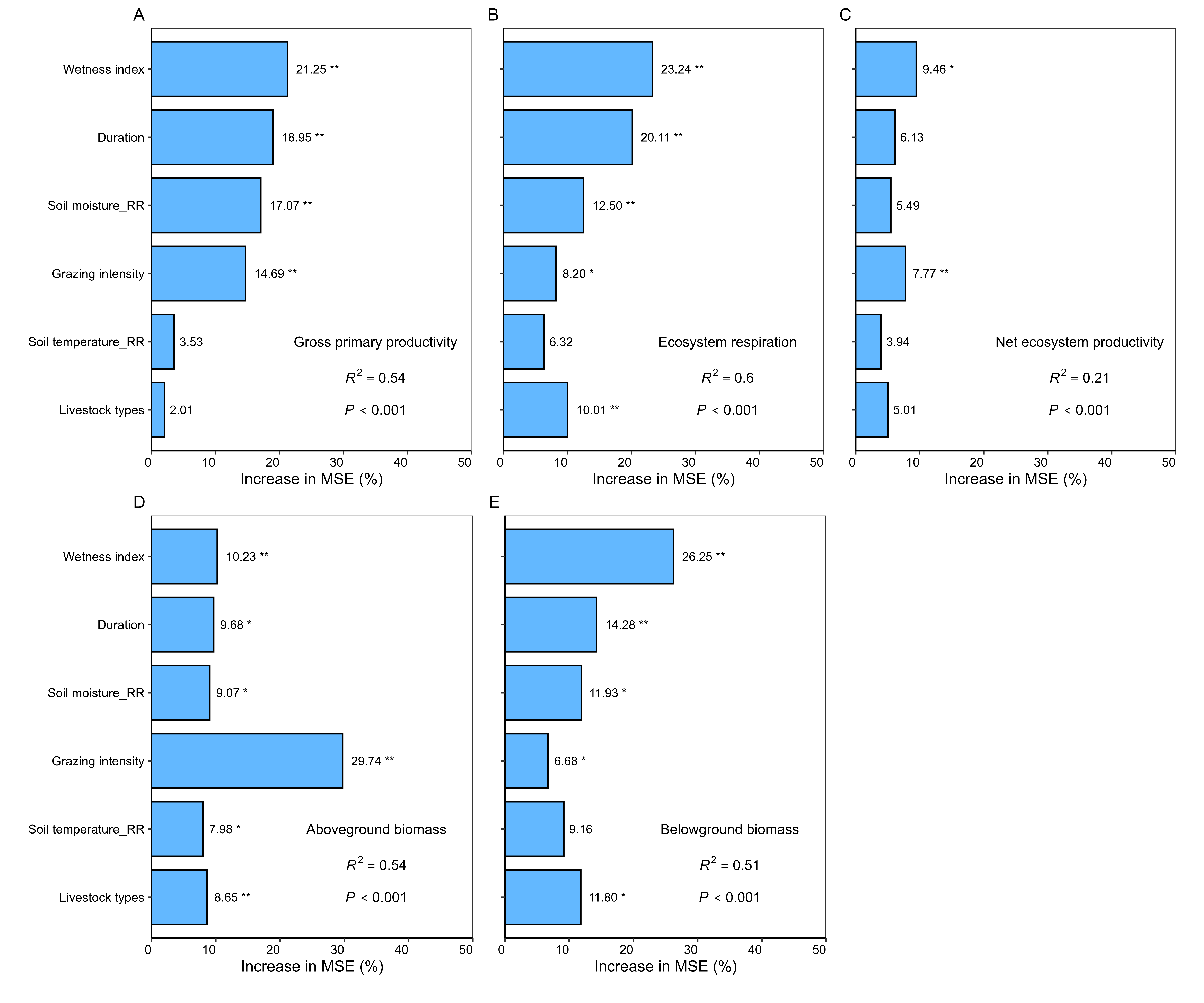

**Fig.** S14 Relationships between the responses of the ecosystem CO_2_ fluxes and aboveground biomass to grazing based on the field experiment in this study (A-C) and from the meta-analysis (D-F). The marginal (m) and conditional (c) *R*^2^ indicate the proportion of variance explained by fixed effects and by both fixed and random effects (Study), respectively. The regression lines and 95% confidence intervals are shown with lines (solid for significant) and shaded areas, respectively. The sizes of points are proportional to their corresponding weights. RR, response ratio; overall, overall grazing; LG, light grazing; MG, moderate grazing; HG, heavy grazing.

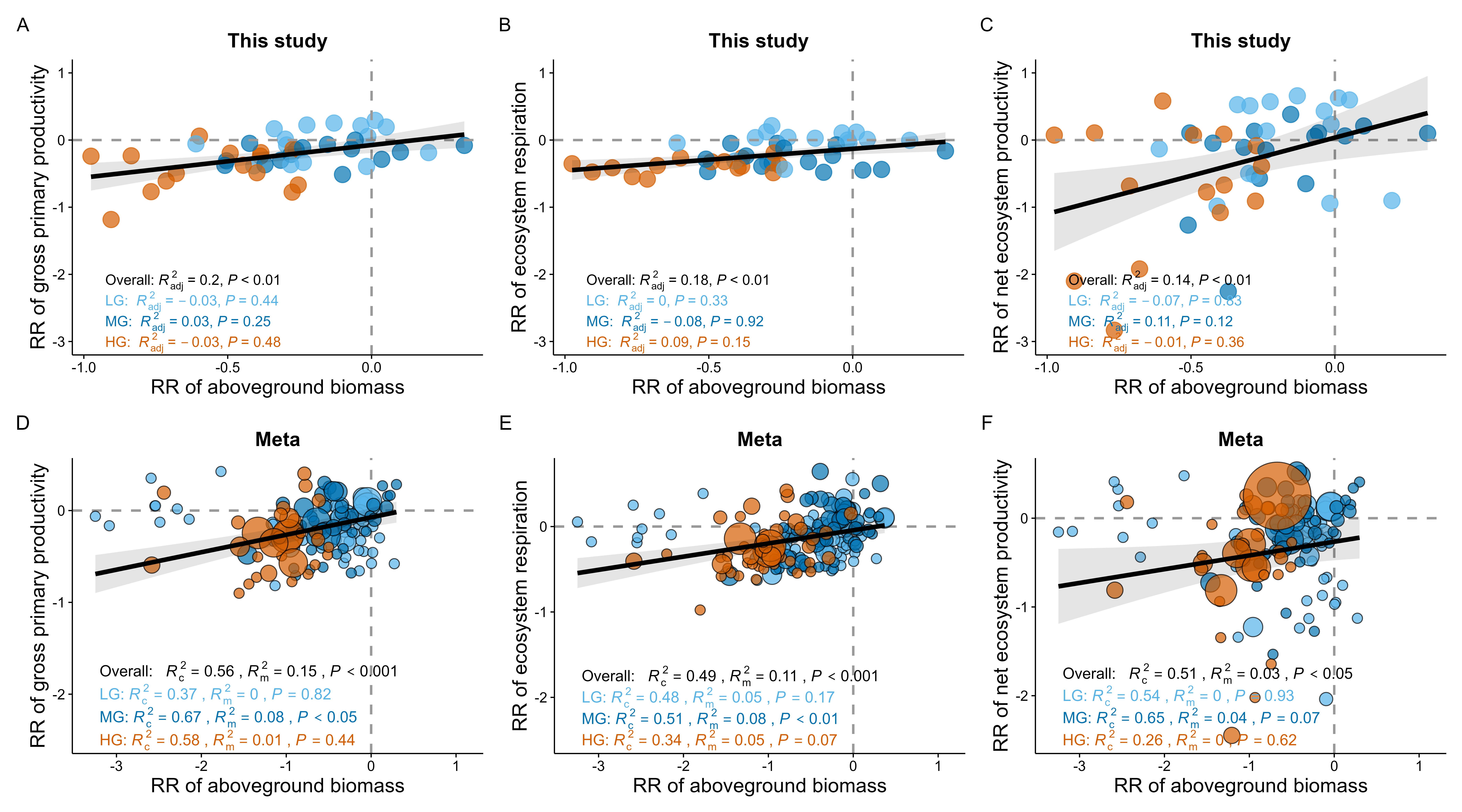

**Fig.** S15 Schematic summary of the effects of grazing on ecosystem carbon dioxide (CO_2_) fluxes, biomass and soil microclimate in the typical steppe of this study and in global grasslands of the meta-analysis. Asterisks (^*^) indicate significant effects on variables at *P* < 0.05. ↑, positive
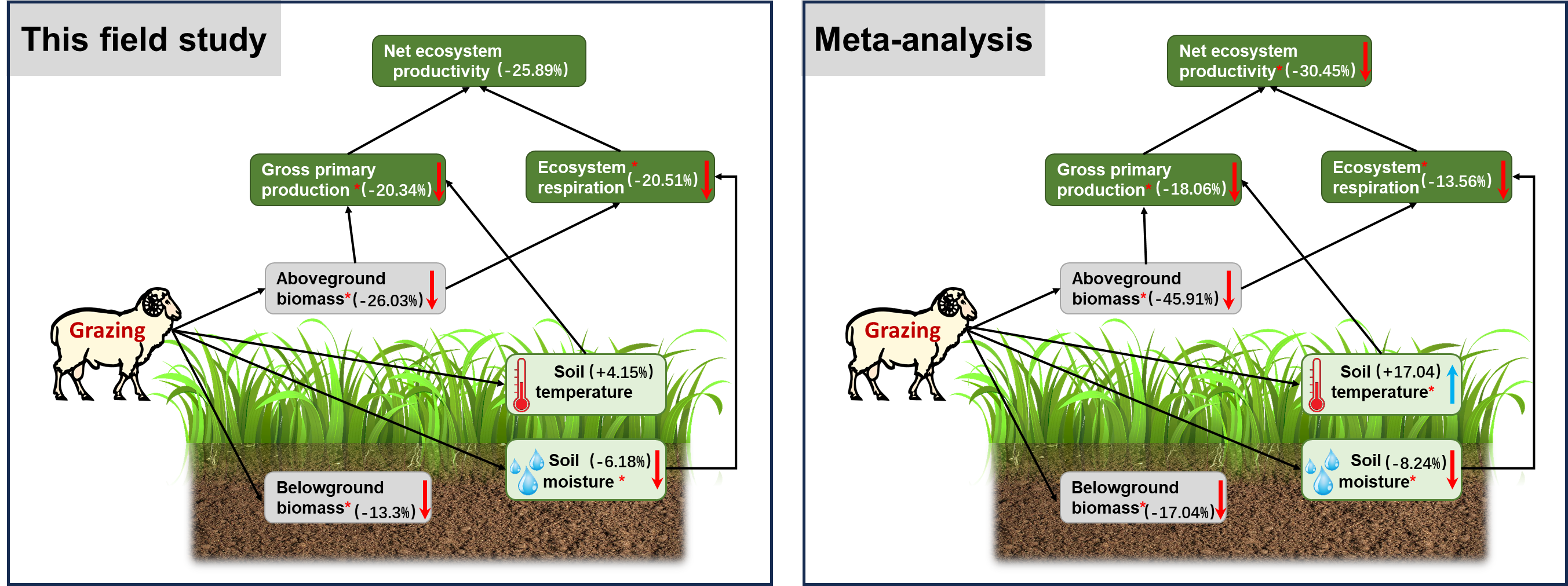
effect response to grazing (blue arrows); ↓, negative effect response to grazing (red arrows).

### **References**

Jiang, Z., Hu, Z., Lai, D., Han, D., Wang, M., Liu, M., Zhang, M., & Guo, M. (2020). Light grazing facilitates carbon accumulation in subsoil in Chinese grasslands: A meta-analysis. *Global Change Biology*, 26(12), 7186–7197. https://doi.org/10.1111/gcb.15326

Shi, R., Su, P., Zhou, Z., Yang, J., & Ding, X. (2022). Comparison of eddy covariance and automatic chamber‐based methods for measuring carbon flux. *Agronomy Journal*, 114. <https://doi.org/10.1002/agj2.21031>

Wan, L., Liu, G., & Su, X. (2025). Global meta-analysis reveals different grazing management strategies change greenhouse gas emissions and global warming potential in grasslands. Geography and Sustainability, 6(3), 100251. https://doi.org/10.1016/j.geosus.2024.09.012

Zhang, R., Tian, D., Chen, H. Y. H., Seabloom, E. W., Han, G., Wang, S., Yu, G., Li, Z., & Niu, S. (2022). Biodiversity alleviates the decrease of grassland multifunctionality under grazing disturbance: A global meta-analysis. *Global Ecology and Biogeography*, 31(1), 155–167. https://doi.org/10.1111/geb.13408

Zhou, G., Luo, Q., Chen, Y., Hu, J., He, M., Gao, J., Zhou, L., Liu, H., & Zhou, X. (2019). Interactive effects of grazing and global change factors on soil and ecosystem respiration in grassland ecosystems: A global synthesis. *Journal of Applied Ecology*, *56*(8), 2007–2019. https://doi.org/10.1111/1365-2664.13443.

### **Reference list for meta-analysis:**

1. Bagheri Shirvan, M., Dijkstra, F. A., & Gonzalez, L. A. (2025). Short-term effect of grazing on net ecosystem exchange and fluxes of greenhouse gases in C_3_ and C_4_ pastures during the growing season. *Agriculture, Ecosystems & Environment*, *383,* 109538.
2. Chang, Q., Xu, T., Ding, S., Wang, L., Liu, J., Wang, D., Wang, Y., Li, Z., Zhao, X., Song, X., & Pan, D. (2020). Herbivore assemblage as an important factor modulating grazing effects on ecosystem carbon fluxes in a meadow steppe in Northeast China. *Journal of Geophysical Research: Biogeosciences*, *125*(9), e2020JG005652.
3. Chen, J., Luo, Y., Xia, J., Zhou, X., Niu, S., Shelton, S., Guo, W., Liu, S., Dai, W., & Cao, J. (2018). Divergent responses of ecosystem respiration components to livestock exclusion on the Qinghai Tibetan Plateau. *Land Degradation & Development*, *29*(6), 1726–1737.
4. Chen, J., Shi, W., & Cao, J. (2015). Effects of grazing on ecosystem CO_2_ exchange in a meadow grassland on the Tibetan Plateau during the growing season. *Environmental Management*, *55*(2), 347–359.
5. Chimner, R. A., & Welker, J. M. (2011). Influence of grazing and precipitation on ecosystem carbon cycling in a mixed-grass prairie. *Pastoralism: Research, Policy and Practice*, *1*(1), 20.
6. Du, C., Zhou, G., & Gao, Y. (2022). Grazing exclusion alters carbon flux of alpine meadow in the Tibetan Plateau. *Agricultural and Forest Meteorology*, *314*, 108774.
7. Gao J J. (2014). Effects of wind erosion and grazing on ecosystem carbon cycle in temperate steppe. *Henan University* (in Chinese).
8. Han, D., Zhu, X., Yu, P., Hu, Y., Jia, H., & Li, D. (2017). Effects of long-term fencing on ecosystem carbon exchange in a meadow steppe in Central Asia. *Fresenius Environmental Bulletin*, *26*, 6421–6431.
9. Hou, L., Liu, Y., Du, J., Wang, M., Wang, H., & Mao, P. (2016). Grazing effects on ecosystem CO_2_ fluxes differ among temperate steppe types in Eurasia. *Scientific Reports*, *6*(1), 29028.
10. Hu Y., Zhu X., Jia H., Han D., Hu B., & Li D. (2018). Effects of fencing on ecosystem carbon exchange at meadow steppe in the northern slope of the Tianshan Mountains. *Chinese Journal of Plant Ecology*, *42*(3), 372–381.
11. Hu Y. (2010). Studies of effects of climate change and grazing on greenhouse gases fluxes on the alpine meadow of the Qinghai-Tibetan plateau. *University of Chinese Academy of Sciences* (in Chinese).
12. Hu Y. (2023). Effects of different grazing intensities on carbon fluxes in semi-arid grassland ecosystems. *Shanxi Agricultural University* (In Chinese).
13. Ju, X., Wang, B., Wu, L., Zhang, X., Wu, Q., & Han, G. (2024). Grazing decreases net ecosystem carbon exchange by decreasing shrub and semi-shrub biomass in a desert steppe. *Ecology and Evolution*, 14(6), e11528.
14. Jin, Y.X., Liu, F., Zhang, J., Han, M.Q., Wang, Z.W., Qu, Z.Q. & Han, G.D. (2018) Net ecosystem carbon exchange characteristics in *Stipa breviflora* desert steppe with different stocking rates. *Chinese Journal of Plant Ecology*, 42, 361–371.
15. K’Otuto, G. O., Otieno, D. O., Seo, B., Ogindo, H. O., & Onyango, J. C. (2013). Carbon dioxide exchange and biomass productivity of the herbaceous layer of a managed tropical humid savanna ecosystem in western Kenya. *Journal of Plant Ecology*, *6*(4), 286–297.
16. Lecain, D. R., Morgan, J. A., Schuman, G. E., Reeder, J. D., & Hart, R. H. (2000). Carbon exchange rates in grazed and ungrazed pastures of Wyoming. *Journal of Range Management*, *53*(2), 199–206.
17. LeCain, D. R., Morgan, J. A., Schuman, G. E., Reeder, J. D., & Hart, R. H. (2002). Carbon exchange and species composition of grazed pastures and exclosures in the shortgrass steppe of Colorado. *Agriculture, Ecosystems & Environment*, *93*(1), 421–435.
18. Li B. (2014). Impact of grazing intensities on carbon budget of alpine grassland in Qinghai Lake region. *University of Chinese Academy of Sciences* (in Chinese).
19. Li G. (2015). Effects of fencing and grazing on biomass and the exchange of carbon dioxi of *Leymus* *secalinus* communities. *Shanxi Agricultural University* (In Chinese).
20. Li H., Zhang F., Mao S., Zhu J., He H., Wei Y., Yang Y., & Li Y. (2019). Effects of grazing density on ecosystem CO_2_ exchange of Haibei Alpine Kobresia humilis meadow in Qinghai. *Chinese Journal of Grassland*, *41*(2), 16–21. (In Chinese)
21. Li X. (2025). Effects of different grazing intensities on carbon use efficiency of *Leymus secalinus* grassland ecosystem in Northern Shanxi. *Shanxi Agricultural University* (In Chinese).
22. Liebig, M. A., Kronberg, S. L., Hendrickson, J. R., Dong, X., & Gross, J. R. (2013). Carbon dioxide efflux from long-term grazing management systems in a semiarid region. *Agriculture, Ecosystems & Environment*, *164*, 137–144.
23. Lin, X., Zhang, Z., Wang, S., Hu, Y., Xu, G., Luo, C., Chang, X., Duan, J., Lin, Q., Xu, B., Wang, Y., Zhao, X., & Xie, Z. (2011). Response of ecosystem respiration to warming and grazing during the growing seasons in the alpine meadow on the Tibetan plateau. *Agricultural and Forest Meteorology*, *151*(7), 792–802.
24. Liu, Y., Tenzintarchen, Geng, X., Wei, D., Dai, D., & Xu-Ri. (2020). Grazing exclusion enhanced net ecosystem carbon uptake but decreased plant nutrient content in an alpine steppe. *Catena*, *195*, 104799.
25. Liu Y. (2018). Greenhouse gas emissions of alpine meadow grazing systems on the Qinghai-Tibet Plateau. *Lanzhou University* (In Chinese).
26. Lu Q. (2025). Regulatory mechanisms of seasonal grazing on key processes of soil carbon transformation in desert steppe. *Ningxia University* (In Chinese).
27. Luo, C., Bao, X., Wang, S., Zhu, X., Cui, S., Zhang, Z., Xu, B., Niu, H., Zhao, L., & Zhao, X. (2015). Impacts of seasonal grazing on net ecosystem carbon exchange in alpine meadow on the Tibetan Plateau. *Plant and Soil*, *396*(1), 381–395.
28. Luo, C., Wang, S., Zhang, L., Wilkes, A., Zhao, L., Zhao, X., Xu, S., & Xu, B. (2020). CO_2_, CH_4_ and N_2_O fluxes in an alpine meadow on the Tibetan Plateau as affected by N-addition and grazing exclusion. *Nutrient Cycling in Agroecosystems*, *117*(1), 29–42.
29. Mei, B., Yue, H., Zheng, X., McDowell, W. H., Zhao, Q., Zhou, Z., & Yao, Z. (2019). Effects of grazing pattern on ecosystem respiration and methane flux in a sown pasture in Inner Mongolia, China. *Atmosphere*, *10*(1), 5.
30. Munjonji, L., Ayisi, K. K., Mudongo, E. I., Mafeo, T. P., Behn, K., Mokoka, M. V., & Linstädter, A. (2020). Disentangling Drought and Grazing Effects on Soil Carbon Stocks and CO_2_ Fluxes in a Semi-Arid African Savanna. *Frontiers in Environmental Science*, *8*.
31. Nakano, T., Bavuudorj, G., Iijima, Y., & Ito, T. Y. (2020). Quantitative evaluation of grazing effect on nomadically grazed grassland ecosystems by using time-lapse cameras. *Agriculture, Ecosystems & Environment*, *287*, 106685.
32. Ondier, J. O., Okach, D. O., Onyango, J. C., & Otieno, D. O. (2021). Ecosystem productivity and CO_2_ exchange response to the interaction of livestock grazing and rainfall manipulation in a Kenyan savanna. *Environmental and Sustainability Indicators*, *9*, 100095.
33. Qi, Y., Wei, D., Wang, Z., Zhao, H., Fan, J., Tao, J., & Wang, X. (2024). Optimizing restoration duration to maximize CO_2_ uptake on the Tibetan Plateau. *Catena*, *241*.
34. Qiao C. (2011). Alpine meadow ecosystem carbon budget under different grazing densities in growing season. *University of Chinese Academy of Sciences* (In Chinese).
35. Qiao J., Ding R., Zhu G., & Wang C. (2018). Effects of mixed grazing on soil greenhouse gas fluxes. *Journal of Northern Agriculture*, *46*(4), 105–109.
36. Qiao L. (2014). Effects of water and nitrogen additions on ecosystem carbon exchanges in a grazing ecosystem of Inner Mongolia typical steppe. *Inner Mongolia University* (In Chinese).
37. Rong, Y., Johnson, D. A., Wang, Z., & Zhu, L. (2017). Grazing effects on ecosystem CO_2_ fluxes regulated by interannual climate fluctuation in a temperate grassland steppe in northern China. *Agriculture, Ecosystems & Environment*, *237*, 194–202.
38. Sagar, R., Li, G. Y., Singh, J. S., & Wan, S. (2019). Carbon fluxes and species diversity in grazed and fenced typical steppe grassland of Inner Mongolia, China. *Journal of Plant Ecology*, *12*(1), 10.
39. Sharkhuu, A., Plante, A. F., Enkhmandal, O., Gonneau, C., Casper, B. B., Boldgiv, B., & Petraitis, P. S. (2016). Soil and ecosystem respiration responses to grazing, watering and experimental warming chamber treatments across topographical gradients in northern Mongolia. *Geoderma*, *269*, 91–98.
40. Tan X. (2017). Effects of grazing on productivity and ecosystem carbon fluxes in a semiarid grassland, Inner Mongolia. *University of Chinese Academy of Sciences* (In Chinese).
41. Tang Y., Wu Y., Wu K., Guo Z., Liang C., Wang M., & Chang P. (2019). Changes in trade-offs of grassland ecosystem services and functions under different grazing intensities. *Chinese Journal of Plant Ecology*, *43*(5), 408–417.
42. Wachiye, S., Pellikka, P., Rinne, J., Heiskanen, J., Abwanda, S., & Merbold, L. (2022). Effects of livestock and wildlife grazing intensity on soil carbon dioxide flux in the savanna grassland of Kenya. *Agriculture, Ecosystems & Environment*, *325*, 107713.
43. Wang B. (2023). Effects of Different Grazing Intensities on Ecosystem Carbon Exchange in *Stipa breviflora* Desert Steppe. *Inner Mongolia Agricultural University* (In Chinese).
44. Wang, D., Wu, G.-L., Liu, Y., Yang, Z., & Hao, H.-M. (2015). Effects of grazing exclusion on CO_2_ fluxes in a steppe grassland on the Loess Plateau (China). *Ecological Engineering*, *83*, 169–175.
45. Wang, Y., Zhao, Q., Wang, Z., Zhao, M., & Han, G. (2023). Overgrazing leads to decoupling of precipitation patterns and ecosystem carbon exchange in the desert steppe through changing community composition. *Plant and Soil*, 486(1–2), 607–620.
46. Wang X. (2023). Effects of grazing on the key processes of carbon cycle in alpine grassland—A case study in Haibei alpine meadow, Qinghai Province*.* *Lanzhou University* (In Chinese).
47. Wang Y. (2017). Responses of ecosystem CO_2_ exchange to nitrogen addition, water addition and grazing in Songnen meadow steppe. *Northeast Normal University* (In Chinese).
48. Wen, C., Huang, J., Shan, Y., Yang, D., Mu, L., Zhang, P., Liu, X., Chang, H., & Ye, R. (2025). Effects of Nitrogen and Water Addition on Ecosystem Carbon Fluxes in a Grazing Desert Steppe. *Agronomy-Basel*, 15(8), 2016.
49. Welker, J. M., Fahnestock, J. T., Povirk, K. L., Bilbrough, C. J., & Piper, R. E. (2004). Alpine grassland CO_2_ exchange and nitrogen cycling: Grazing history effects, medicine bow range, Wyoming, USA. *Arctic Antarctic and Alpine Research*, *36*(1), 11–20.
50. Wilsey, B. J., Parent, G., Roulet, N. T., Moore, T. R., & Potvin, C. (2002). Tropical pasture carbon cycling: Relationships between C source/sink strength, above-ground biomass and grazing. *Ecology Letters*, *5*(3), 367–376.
51. Xing P., Li G., Chen X., LI D., Wang C., Dong K., & Zhao X. (2019). Effects of grazing on carbon exchange in a *Leymus secalinus* grassland ecosystem in the agro-pastoral ecotone of Northern Shanxi. *Acta Prataculturae Sinica*, *28*(10), 1–11. (In Chinese)
52. Xu, G., Kang, X., Li, W., Li, Y., Chai, Y., Wu, S., Zhang, X., Yan, Z., Kang, E., Yang, A., Niu, Y., Wang, X., & Yan, L. (2022). Different grassland managements significantly change carbon fluxes in an alpine meadow. *Frontiers in Plant Science*, *13*.
53. Yan, R., Zhang, Y., Wang, M., Li, R., Jin, D., Xin, X., & Li, L. (2021). Interannual variation in ecosystem respiration in an Inner Mongolian meadow steppe in response to livestock grazing. *Ecological Indicators*, *131*, 108121.
54. Yan W., Sun G., Zhang C., He J., & Zhang N. (2018). Impacts of experimental warming and moderate grazing on ecosystem carbon exchange and its compositions in an alpine meadow on the eastern Qinghai-Tibetan Plateau. *Chinese Journal of Applied & Environmental Biology*, *24*(1), 132–139. (In Chinese)
55. Yang, Z. (2014). Effects of Grazing management regimes on carbon flux of a semiarid grassland in Inner Mongonia. *University of Chinese Academy of Sciences* (In Chinese).
56. Yu, H., Wang, X., Wu, Y., Wang, C., Yan, R., Xu, D., Yan, Y., & Xin, X. (2025). Light grazing tends to enhance ecosystem carbon sequestration and resource use efficiency in a meadow steppe of Northern China. *Agricultural and Forest Meteorology*, *372*, 110690.
57. Zhao, J., Luo, T., Li, R., Li, X., & Tian, L. (2016). Grazing effect on growing season ecosystem respiration and its temperature sensitivity in alpine grasslands along a large altitudinal gradient on the central Tibetan Plateau. *Agricultural and Forest Meteorology*, *218–219*, 114–121.
58. Zhao W. (2025). Effects of four seasons grazing on alpine grassland carbon sequestration and soil microbial communities. *Qinghai University* (In Chinese).
59. Zhou D. (2018). Effects of land use types on greenhouse gas emissions and ecosystem carbon balance in the typical steppe of Inner Mongolia. *Inner Mongolia University* (In Chinese).
60. Zhou P. (2011). Effects of the different grazing intensity on greenhouse gas fluxes of Inner Mongolia grassland. *Inner Mongolia Agricultural University* (In Chinese).
61. Zhou P., Han G., Wang C., Jiang Y., & Tang S. (2011). Effects of stocking rates on carbon flux in the desert grassland ecological system of Inner Mongolia. *Journal of Inner Mongolia Agricultural University*, *32*(4), 59–64 (In Chinese).
62. Zhu, L., Johnson, D. A., Wang, W., Ma, L., & Rong, Y. (2015). Grazing effects on carbon fluxes in a Northern China grassland. *Journal of Arid Environments*, *114*, 41–48.
